# Low-Dimensional Spatio-Temporal Dynamics Underlie Cortex-Wide Neural Activity

**DOI:** 10.1101/2020.01.05.895177

**Authors:** Camden J. MacDowell, Timothy J. Buschman

## Abstract

Cognition arises from the dynamic flow of neural activity through the brain. To capture these dynamics, we used mesoscale calcium imaging to record neural activity across the dorsal cortex of awake mice. We found that the large majority of variance in cortex-wide activity (∼75%) could be explained by a limited set of ∼14 ‘motifs’ of neural activity. Each motif captured a unique spatio-temporal pattern of neural activity across the cortex. These motifs generalized across animals and were seen in multiple behavioral environments. Motif expression differed across behavioral states and specific motifs were engaged by sensory processing, suggesting the motifs reflect core cortical computations. Together, our results show that cortex-wide neural activity is highly dynamic, but that these dynamics are restricted to a low-dimensional set of motifs, potentially to allow for efficient control of behavior.

## Introduction

The brain is a complex, interconnected network of neurons. Neural activity flows through this network, carrying and transforming information to support behavior. Previous work has associated particular computations with specific spatio-temporal patterns of neural activity across the brain (Buschman and Miller, 2007; Stringer et al., 2019a; Hutchison et al., 2013). For example, sequential activation of primary sensory and then higher-order cortical regions underlies perceptual decision making in both mice (Guo et al., 2014) and monkeys (Romo and de Lafuente, 2013; Siegel et al., 2011). Similarly, specific spatio-temporal patterns of cortical regions are engaged during goal-directed behaviors (Allen et al., 2017), motor planning (Chen et al., 2017), evidence accumulation (Pinto et al., 2019), motor learning (Makino et al., 2017), and sensory processing (Mohajerani et al., 2013). Previous work has begun to codify these dynamics, either in the synchronous activation of brain regions (Fries, 2015; Hutchison et al., 2013) or in the propagation of waves of neural activity within and across cortical regions (Muller et al., 2018; Zanos et al., 2015). Together, this work suggests cortical activity is highly dynamic, evolving over both time and space, and that these dynamics play a computational role in cognition (Buonomano and Maass, 2009; Miller and Wilson, 2008).

However, despite this work, the nature of cortical dynamics is still not well understood. Previous work has been restricted to specific regions and/or specific behavioral states and so, we do not yet know how neural activity evolves across the entire cortex, whether dynamics are similar across individuals, or how dynamics relate to behavior. This is due, in part, to the difficulty of quantifying the spatio-temporal dynamics of neural activity across the brain.

To address this, we used mesoscale imaging to measure neural activity across the dorsal cortical surface of the mouse brain (Silasi et al., 2016). Then, using a convolutional factorization approach, we identified dynamic ‘motifs’ of cortex-wide neural activity. Each motif captured a unique spatio-temporal pattern of neural activity as it evolved across the cortex. Importantly, because motifs captured the dynamic flow of neural activity across regions, they explained cortex-wide neural activity better than ‘functional connectivity’ network measures.

Surprisingly, the motifs clustered into a limited set of ∼14 different spatio-temporal ‘basis’ motifs that were consistent across all animals. The basis motifs captured the majority of the variance in neural activity in different behavioral states and in multiple sensory and social environments. Specific motifs were selectively engaged by each environment and by sensory stimuli, suggesting the motifs reflect core cortical computations, such as visual or tactile processing. Together, our results suggest cortex-wide neural activity is highly dynamic but that these dynamics are low-dimensional: they are constrained to a small set of possible spatio-temporal patterns.

## Results

### Discovery of spatio-temporal motifs of cortical activity in awake, head-fixed mice

We performed widefield ‘mesoscale’ calcium imaging of the dorsal cerebral cortex of awake, head-fixed mice expressing the fluorescent calcium indicator GCaMP6f in cortical pyramidal neurons (Fig. 1A; see Methods for details, Chen et al., 2013). A translucent-skull prep provided optical access to dorsal cortex, allowing us to track the dynamic evolution of neural activity across multiple brain regions, including visual, somatosensory, retrosplenial, parietal, and motor cortex (Fig. 1A, inset and Fig. 1 supplement 1, Silasi et al., 2016). We initially characterized the dynamics of ‘spontaneous’ neural activity when mice were not explicitly performing a specific behavior (Fig. 1B, N=48 sessions across 9 mice, 5-6 sessions per mouse, each session lasted 12 minutes, yielding a total of 9.6 hours of imaging). These recordings revealed rich dynamics in the spatio-temporal patterns of neural activity across the cortex (Supplemental Movie 1, as also seen by Cramer et al., 2019; Shimaoka et al., 2019; Murphy et al., 2016; Mohajerani et al., 2013)

**Figure 1.**
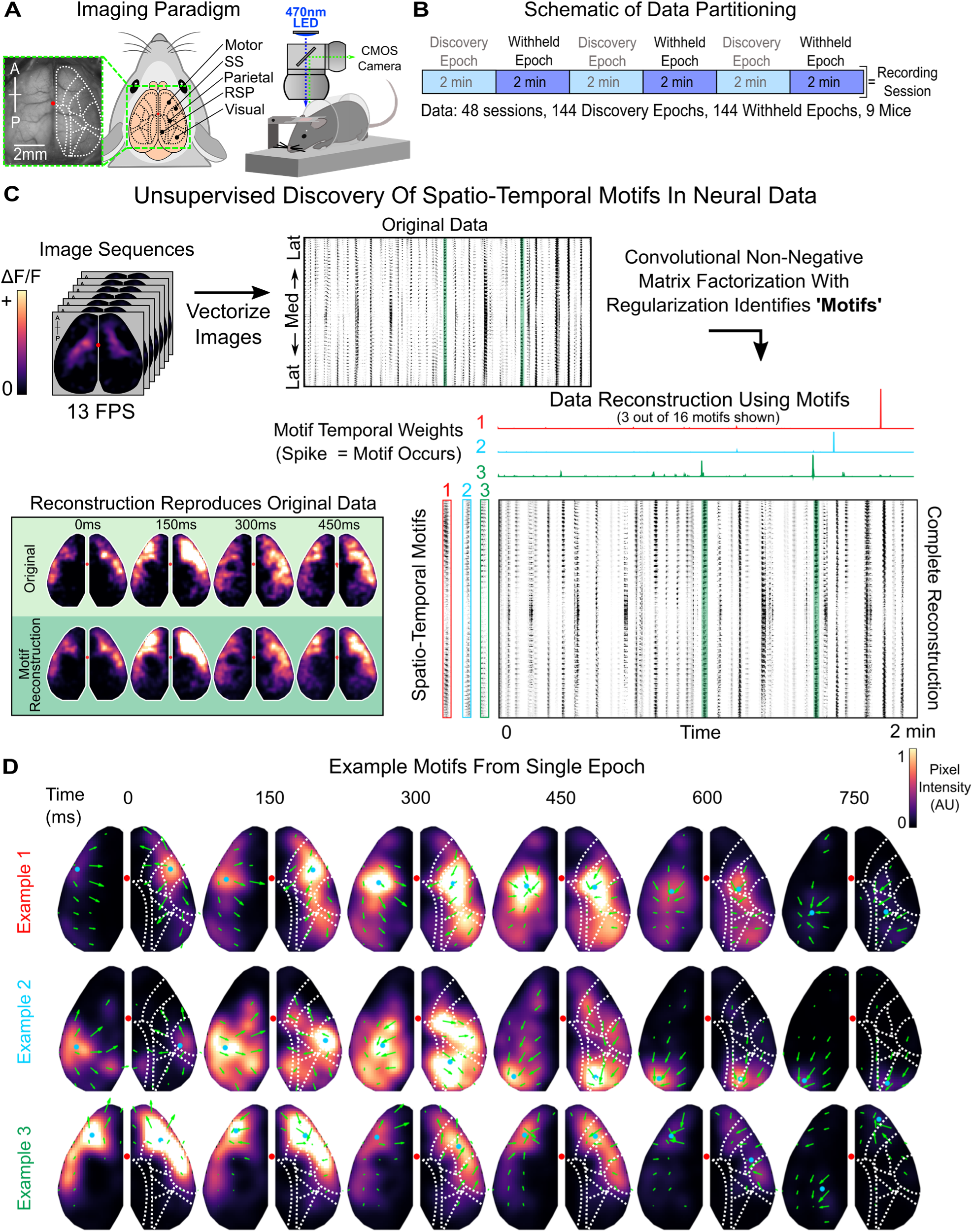
Discovery of spatio-temporal patterns in cortical activity of awake, head-fixed mice. **(A)** Schematic of imaging paradigm. Mice expressing GCaMP6f in cortical pyramidal neurons underwent a translucent skull prep to allow mesoscale imaging of neural activity across the majority of dorsal cortex. Red dot denotes bregma. Cortical parcellation follows Allen Brain Atlas. General anatomical parcels are labeled. Motor, motor cortex; SS, somatosensory cortex; Parietal, parietal cortex; RSP, retrosplenial cortex; Visual: visual cortex (See Fig. 1 supplement 1 for complete parcellation of 24 regions; 12 per hemisphere). **(B)** Schematic of data partitioning. 9 mice were imaged for 12 minutes a day for 5-6 consecutive days. Recording sessions (N=48) were divided into 2-minute epochs (N=144). Alternating epochs were used for discovering spatio-temporal motifs in neural activity or were withheld for testing generalization of motifs. **(C)** Schematic of unsupervised discovery of spatio-temporal motifs from a single epoch. Mesoscale calcium imaging captured patterns of neural activity, measured by change in fluorescence (ΔF/F), across the dorsal cortex (top-left). Movies were vectorized (top-middle, black=activity) and then decomposed into dynamic, spatio-temporal motifs (bottom-right; 3 example motifs shown along left, temporal weightings along top). Convolving motifs with temporal weightings reconstructed the original movie (bottom-left; snapshot of data is highlighted in green throughout, corresponding to activity in motif 3, which is shown in image format in bottom row of **D**). Note: only 3 out of 16 example motifs and their corresponding temporal weightings are shown; data reconstruction in bottom right used all 16 motifs. **(D)** Timecourses of the 3 example motifs in panel **C** showing spatio-temporal patterns of neural activity across dorsal cortex. Arrows indicate direction of flow of activity across subsequent timepoints (see Methods for details). Blue dot denotes center of mass of the most active pixels in each hemisphere (>=95% intensity). Red dot denotes bregma. Dotted white lines outline anatomical parcels as in **A**. Only every other timepoint is shown. For visualization, motifs were filtered with 3D gaussian (across space and time), and intensity scale is normalized for each motif. Intensity value is arbitrary as responses are convolved with independently scaled temporal weightings to reconstruct the normalized ΔF/F fluorescence trace (see Methods for details).

Our goal was to capture, quantify, and characterize the dynamic patterns of activity in an unbiased manner. To do so, we used convolutional non-negative matrix factorization (CNMF, Mackevicius et al., 2019) to discover repeated spatio-temporal patterns in the neural activity, in an unsupervised way (Fig. 1C; see Methods and Fig. 1 supplement 2 for details, including robustness to parameters). CNMF identified ‘motifs’ of neural activity; these are dynamic patterns of neural activity that extend over space and time (Fig. 1C, bottom left, shows an example motif and the corresponding original data). Once identified, the algorithm uses these motifs to reconstruct the original dataset by temporally weighting them across the entire recording session (Fig. 1C; transients in temporal weightings indicate motif expression, see Supplemental Movie 2 for comparison of reconstruction to original data). Importantly, overlapping temporal weightings between motifs are penalized, which biases factorization towards only one motif being active at a given point in time. This allows us to capture the spatio-temporal dynamics of neural activity as a whole, rather than decomposing activity into separate spatial and temporal parts (see Methods for details).

Figure 1D shows three example motifs identified by CNMF from a single 2-minute recording epoch. Many of the identified motifs show dynamic neural activity that involves the sequential activation of multiple regions of cortex (top two rows in Fig. 1D). For example, example motif 1 starts in somatosensory/motor regions and, over the course of a few hundred milliseconds, propagates posteriorly before ending in the parietal and visual cortices (Fig. 1D, top row). To aid in visualizing these dynamics, the arrows overlaid on Figure 1D show the direction and magnitude of activity propagation of the top 50% most active pixels between subsequent timepoints (as in Afrashteh et al., 2017; see Methods for details). Other motifs were more spatially restricted, engaging either one (or more) brain regions simultaneously (e.g. third row in Fig. 1D). In total, we identified 2622 motifs across 144 different 2-minute epochs of imaging (3 independent epochs from each of 48, 12-minute recording sessions; Fig. 1B, light blue ‘discovery epochs’).

Motifs captured the flow of activity across the brain during a brief time period (∼1 second). By tiling different motifs across time, the entire 2-minute recording epoch could be reconstructed. On average, each 2-minute recording epoch could be reconstructed by combining ∼19 motifs (Fig 2A. median; 95% Confidence Interval (CI): 18-20). This captured 89.05% of the total variance of neural activity on average (Fig. 2B; 89.05% median explained variance, CI: 87.78-89.68%; N=144 discovery epochs; see Methods for details). To achieve this, individual motifs occurred repeatedly during a recording session. Over half of the motifs occurred at least 3 times during a 2-minute epoch, with motifs occurring 2.48 times per minute on average (Fig. 2C; CI: 2.38-2.59). The broad distribution of the frequency of motifs suggests all motifs are required to explain cortex-wide neural activity. Indeed, the cumulative percent explained variance (PEV) in neural activity captured by individual motifs shows a relatively gradual incline (Fig. 2D, see Methods for details). On average, no single motif captured more than 20% of the variance of the recording epoch, and 14 motifs were needed to capture over 90% of the relative PEV. Importantly, the number of discovered motifs and their explained variance was robust to the changes in the regularization hyperparameter of the CNMF algorithm, suggesting it is a true estimate of the number of motifs needed and not a consequence of our analytical approach (Fig. 1 supplement 2, see Methods for details).

**Figure 2.**
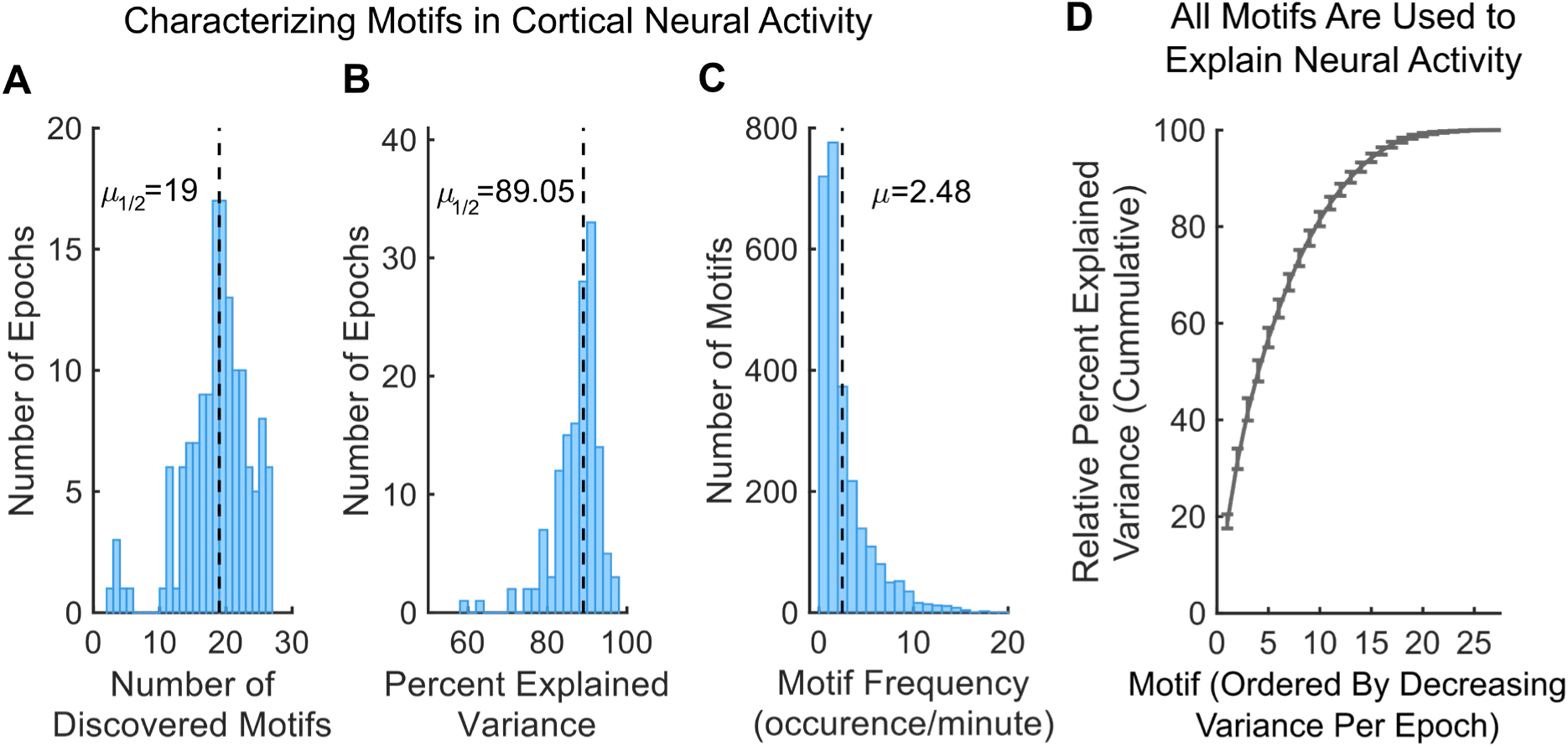
Motifs capture majority of variance in neural activity. **(A)** Distribution of number of discovered motifs per discovery epoch (N=144). Dotted line indicates median. 2622 motifs were discovered in total. **(B)** Distribution of the total percent of variance in neural activity explained by motifs per discovery epoch. Dotted line indicates median. **(C)** Distribution of how often motifs occurred during discovery epochs. A motif was considered active when its temporal weighting was 1 standard deviation above its mean (e.g. transients in Fig. 1C in occurrences per minute; see Methods for details). Dotted line indicates mean. **(D)** Cumulative sum of relative percent explained variance (PEV) of each motif in withheld epochs. Relative PEV was calculated as the PEV of each motif divided by the sum of all motif PEVs in an epoch. For each epoch, motifs are ordered by their relative PEV (i.e. the first motif is the most common motif, which is not necessarily the same motif for all epochs). Line and error bars indicate mean and 95% CI, respectively. All p-values estimated with Wilcoxon Signed-Rank tests.

### Motifs capture the dynamic flow of neural activity across the cortex

Next, we tested whether motifs simply reflected the co-activation of brain regions or if they captured the dynamic flow of neural activity between regions. Previous work has found neural activity can be explained by the simultaneous activation of a coherent network of brain regions (i.e. zero-lag, first-order correlations, as seen in functional connectivity analyses; Hutchison et al., 2013). The CNMF approach used here is a generalization of such approaches; it can capture spatio-temporal dynamics in the motifs, but it is not required to do so if dynamics are not necessary to capture variance in neural activity. Therefore, to test whether neural activity is dynamic, we tested whether dynamics were a necessary component of the motifs.

First, we determined whether motifs simply reflected the static engagement of a network of regions. To this end, we measured the autocorrelation of neural activity during the timecourse of each motif. Consistent with dynamic motifs, the correlation of activity patterns within a motif quickly decayed with time (Fig. 3A; mean half-life τ across all motifs was 113ms +/- 2ms, bootstrap; when fit to individual motifs, 25%-50%-75% of τ was 66ms - 117ms - 210ms; see Methods for details). While activity patterns at adjacent motif timepoints (75ms apart) were spatially correlated (Pearson’s r=0.39 CI: 0.38-0.40, p<10^-16^, Wilcoxon Signed-Rank Test versus r=0, right-tailed; N=2622 Motifs), this similarity quickly declined when time points were farther apart (Pearson’s r=0.098 CI: 0.095-0.10 at 375ms and r=0.043, CI: 0.039-0.047 at 600ms; a decrease of 0.29 and 0.35, both p<10^-16^, Wilcoxon Signed-Rank Test). Similarly, the mean spatial pattern of activity of a given motif, averaged across the timecourse of the motif, was dissimilar from individual timepoints within the motif (Fig. 3B; median dissimilarity 0.58, CI: 0.57-0.58 across motifs).

**Figure 3.**
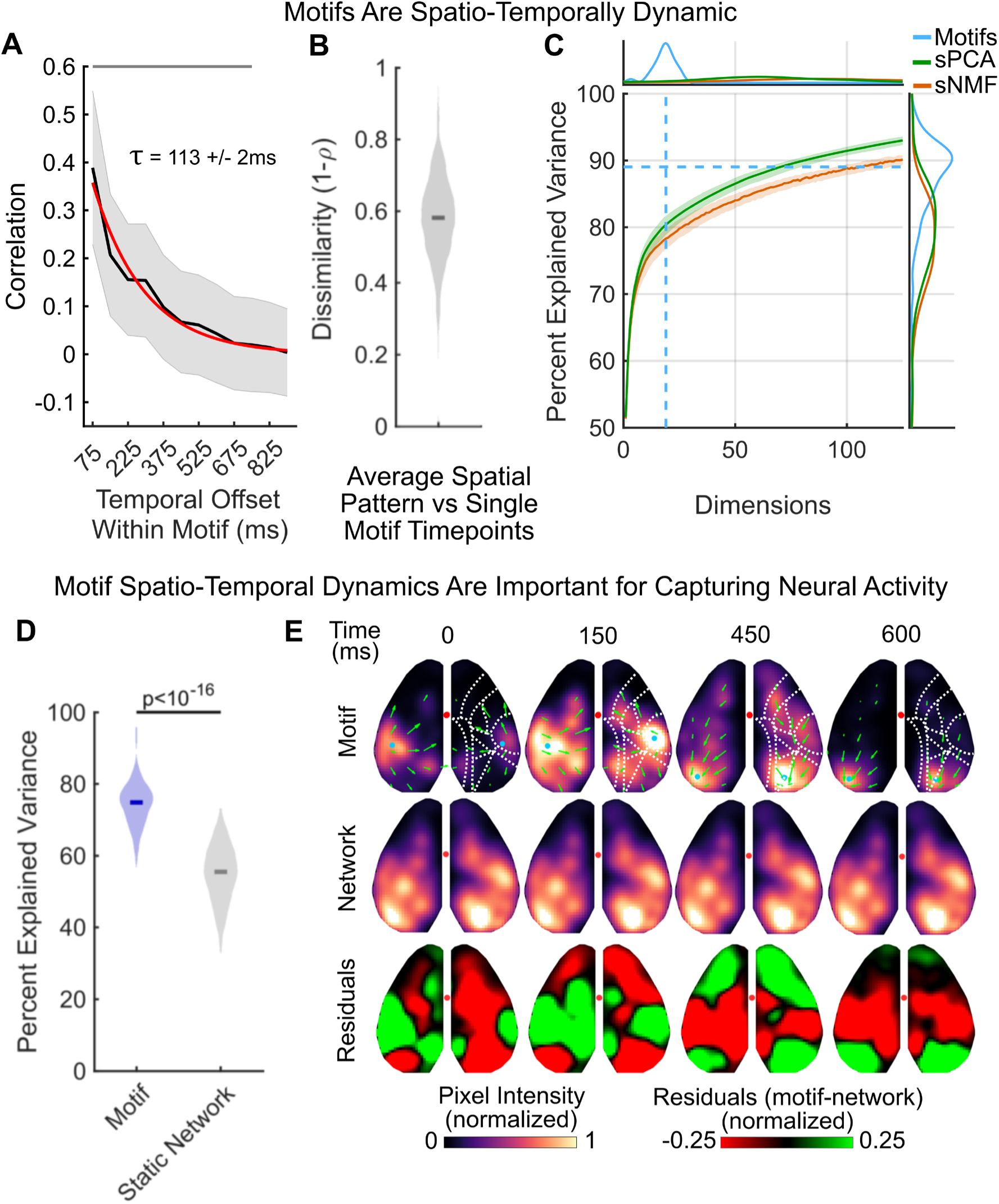
Motifs capture the flow of neural activity across the cortex. **(A)** Cross-temporal autocorrelation of motifs (N=2622). Average spatial correlation of activity (y-axis) was calculated for different temporal offsets (x-axis) within a motif. For example, an offset of 75ms indicates the correlation between timepoint N and timepoints N-1 and N+1 (given sampling frequency of 13.33 Hz). Black line and gray shading denote mean and standard deviation, respectively, across all motifs. Red line shows exponential fit to autocorrelation decay. Mean half-life of autocorrelation decay (τ) across all motifs was 113ms +/- 2ms SEM. **(B)** Average dissimilarity between each timepoint of a dynamic motif and the mean spatial activity of that motif; averaged across frames of the motifs. Full distribution shown, depicts average dissimilarity per motif (N=2622 motifs). Dark line indicates median. **(C)** Comparison of reconstruction of neural data by CNMF motifs (blue), spatial principal components analysis (sPCA, green) and spatial non-negative matrix factorization (sNMF, orange). Central plot shows percent of variance in neural activity explained (y-axis) as a function of number of dimensions included (x-axis). Lines show median variance explained, shaded regions show 95% confidence interval. Dashed light blue lines show median number of motifs discovered (vertical) and the median percent explained variance (horizontal) captured by CNMF motifs across discovery epochs (N=144). **(top)** Plot shows the probability density function (PDF) of number of motifs discovered per discovery epoch (blue), as well as the PDF of the minimum number of dimensions needed to capture the same amount of variance using sPCA (orange) and sNMF (green). **(right)** PDF of percent explained variance by motif reconstructions (blue) across epochs, as well as the PDF of percent of variance explained by sPCA (orange) and sNMF (green) when the number of dimensions is restricted to match the number of discovered motifs in each epoch. For visualization, x-axis is cropped to 125 dimensions. **(D)** Percent of variance in neural activity explained by dynamic motifs (blue) and static networks (grey), defined as the average activity across the motif. Both static networks and motifs are fit to the data in the same manner (see Methods for details). Full distribution shown; dark lines indicate median. Analyses performed on withheld epochs (N=144). **(E)** An example motif (top row; example motif 2 from Fig. 1D) and its corresponding static network (middle row). Bottom row shows normalized residuals between dynamic motif and static network. For calculation of residuals, motif and networks were scaled to the same mean pixel value per timepoint. Display follows Figure 1D. All p-values estimated with Wilcoxon Signed-Rank tests.

Second, we tested whether dynamics were necessary to fit neural data. To do this, we compared the fit of CNMF-derived motifs to alternative decomposition approaches that do not consider temporal dynamics. We used two ‘static’ decomposition techniques that are standards in the field: spatial Principal Components Analysis (sPCA) and spatial Non-Negative Matrix Factorization (sNMF; see Methods for details). Both approaches required >3 times more dimensions to capture the same amount of variance as the motifs (Fig. 3E; on average, 64.5 and 93 dimensions for sPCA and sNMF, respectively). If restricted to 19 dimensions, sPCA and sNMF explained significantly less variance in neural activity than motifs (Fig. 3E; sPCA: 79.87% CI: 78.72-81.46% a difference of 9.18%, p<10^-16^; sNMF 77.95% CI: 76.63-79.42% a difference of 11.10%, p<10^-16^, Wilcoxon Signed-Rank Test). Although there are differences in the number of terms in each dimension, the fact that dynamic motifs can capture significantly more variance than temporally constrained approaches suggests brain activity has complex spatial-temporal dynamics that are not captured by traditional decomposition methods but can be captured by our motifs.

### Motifs generalize to withheld data and across animals

If the neural dynamics captured by CNMF reflect true, repeated, motifs of neural activity, then the motifs identified in one recording session should generalize to other recording sessions. To test if motifs generalized, we refit the motifs identified during a recording ‘discovery’ epoch to withheld data (Fig. 1B, purple, N=144 ‘withheld epochs’). Motifs were fit to new epochs by only optimizing the motif weightings over time (i.e. not changing the motifs themselves, see Methods for details).

Indeed, the motifs generalized; the same motifs could explain 74.82% of the variance in neural activity in withheld data from the same recording session (Fig. 3D, purple, CI: 73.92-76.05%; see Fig. 3 supplement 1A for robustness to sparsity parameter; see Methods for details). This was not just due to fitting activity on average: motifs captured neural activity at each timepoint during a recording epoch, explaining the majority of the variance in the spatial distribution of neural activity in any given frame (60-80%, Fig. 3 supplement 1B).

Dynamics were important for the ability to generalize. To show this, we created ‘static networks’ by averaging neural activity across the timecourse of each motif. This maintained the overall spatial pattern of activity, ensuring the same network of brain regions was activated, but removed any temporal dynamics within a motif (Fig. 3E; see Methods for details). When the static networks were fit to withheld data, they captured significantly less variance in neural activity compared to the dynamic motifs (Fig. 3D, gray; static networks captured 55.50%, CI: 53.74-57.09%; a 19.32% reduction, p<10^-16^, Wilcoxon Signed-Rank Test).

Similarly, motifs generalized across animals: motifs identified in one animal cross-generalized to capture 68.19% of the variance in neural activity in other animals (Fig; 4A, green; CI: 66.74-69.35%, a decrease of 6.63% compared to generalizing within animals, purple, p<10^-16^, Wilcoxon Signed-Rank Test; N=144 withheld epochs; see Methods for details). Together, our results show motifs can generalize across recording session and across animals, suggesting they reflect repeated spatio-temporal dynamics in neural activity.

**Figure 4.**
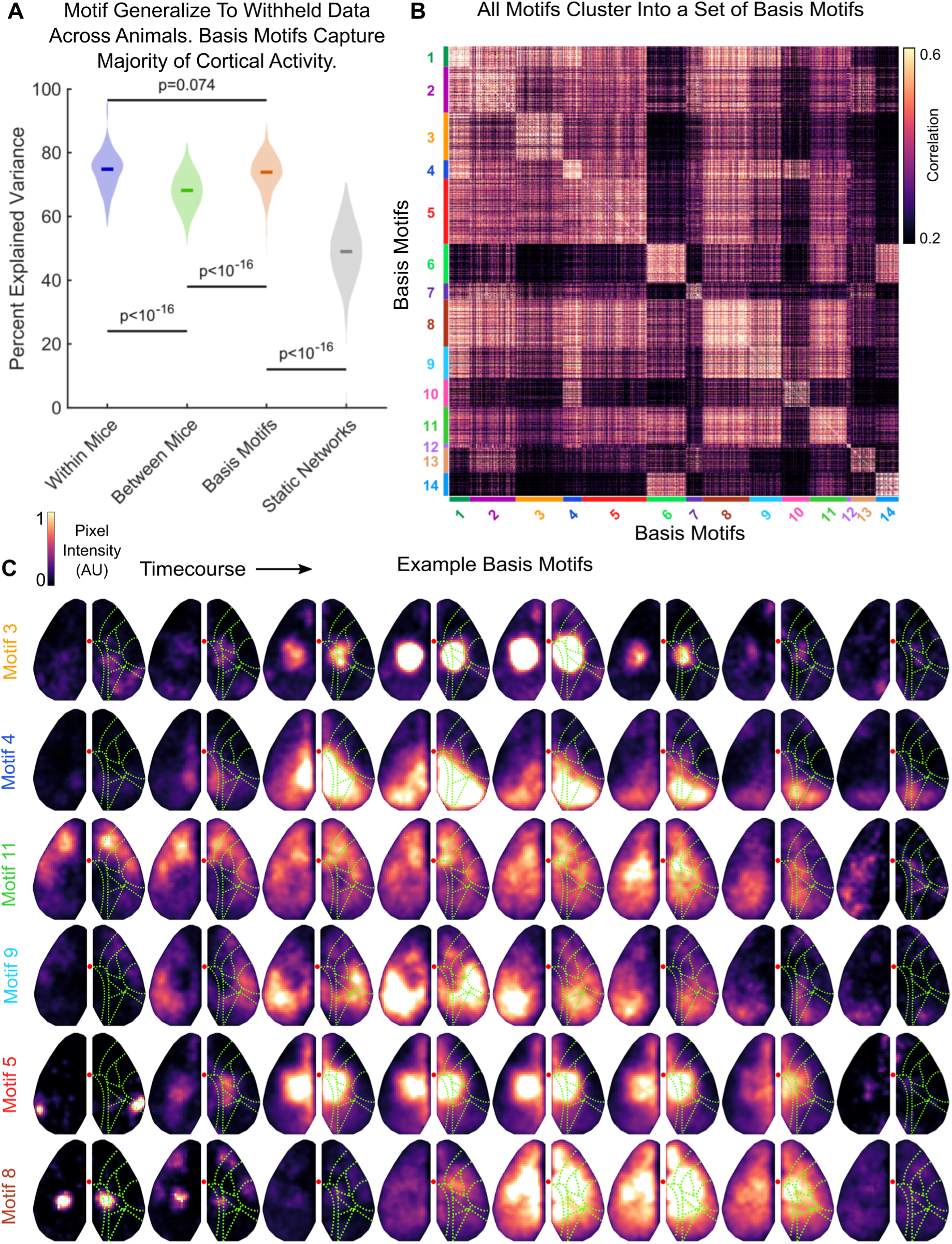
Motifs cluster into a low-dimensional set of basis motifs. **(A)** Comparison of the percent of variance in neural activity explained by motifs from the same mouse (within; purple), by motifs from other mice (between; green), by basis motifs (orange), and by static network versions of basis motifs (gray). Static networks for each basis motifs were derived as in Figure 3 (see Methods for details). All show fit to withheld data (N=144). Full distribution shown; dark lines indicate median. Horizontal lines indicate pairwise comparisons. All p-values estimated with Mann-Whitney U-test. **(B)** Pairwise peak cross-correlation between all 2622 discovered motifs. Motifs are grouped by their membership in basis motif clusters. Basis motif identity is indicated with color code along axes. Group numbering (and thus the sorting of the correlation matrix) is determined by relative variance explained by each basis motif (see Fig. 4 supplement 1A). **(C)** Representative timepoints from example basis motifs. Display follows Fig. 1D, except without direction of flow arrows.

### Motifs cluster into a low-dimensional set of basis motifs

The ability of motifs to generalize across time and animals suggests there may be a set of ‘basis motifs’ that capture canonical patterns of spatio-temporal dynamics. To identify these basis motifs, we used an unsupervised clustering algorithm to cluster all 2622 motifs that were identified across all discovery epochs (clustering done with the Phenograph algorithm using peak of cross-correlation as the distance metric between motifs, Nicosia et al., 2009; Levine et al., 2015, see Methods for details). Motifs clustered into a set of 14 unique groups (Fig. 4B). For each cluster, we defined the basis motif as the mean of the motifs within the ‘core-community’ of each cluster (taken as those motifs with the top 10% most within-cluster nearest neighbors, see Methods for details).

Similar to the motifs discovered within a single session, the basis motifs captured the dynamic engagement of one or more brain regions (Fig. 4C; all basis motifs shown in Supplemental Movie 3). While some basis motifs engaged a single brain region (e.g. motif 3, Fig. 4C), most of the basis motifs captured the propagation of activity across cortex. For example, motif 4 captures the posterior-lateral flow of activity from retrosplenial to visual cortex. Similarly, motif 11 captures a cortex-wide anterior-to-posterior wave of activity that has been previously studied (Greenberg et al., 2018; Matsui et al., 2016; Mitra et al., 2018). As expected, these dynamics were similar to those found in individual recording sessions (e.g. basis motif 9 matches example motif 2 in Fig. 1D).

At the same time, the same brain region, or network of regions, can be engaged in multiple basis motifs. For instance, parietal cortex is engaged in motifs 3, 5 and 8 (Fig. 4C). In motif 3, neural activity remains local to parietal cortex for the duration of the motif. However, in motif 5 parietal activity is prefaced by a burst of activity in rostrolateral cortex. In motif 8, activity starts in parietal areas before spreading across the entire dorsal cortex. Similarly, several motifs (6, 8, 11, and 14) involve coactivation of a network of anterolateral somatosensory and primary motor cortices; a coupling observed in previous mesoscale imaging studies (Musall et al., 2019; Silasi et al., 2016; Vanni et al., 2017). Thus, basis motifs reflect the ordered engagement of multiple brain regions, likely reflecting a specific flow of information through the brain.

Basis motifs explained the large majority of the variance in neural activity across animals (73.91% CI: 73.14-75.19%, Fig. 4A, orange; N=144 withheld epochs). This is about the same amount of variance explained by motifs defined within the same animal (Fig. 4A, purple vs. orange; a 0.91% reduction, p=0.074; Wilcoxon Signed-Rank Test, N=144 withheld epochs). It is significantly more variance explained than when using motifs defined in another animal (Fig. 4A, orange vs. green plots; a 5.72% increase in explained variance; p<10^-16^, Wilcoxon Signed-Rank Test). This improvement is likely because basis motifs are averaged across many instances, removing the spurious noise that exists in individual motifs and resulting in a better estimate of the underlying ‘true’ motif that exist across animals.

As before, dynamics were important for basis motifs; when spatial-temporal dynamics were removed, the variance explained dropped significantly (Fig. 4A, gray vs orange plots; static networks captured 48.99% CI: 47.15-51.30% of variance, 24.92% less than dynamic motifs, p<10^-16^, N=144 withheld epochs, Wilcoxon Signed-Rank Test). Furthermore, all basis motifs were necessary to explain neural activity; the cumulative PEV of motifs followed a gradual rise and no basis motif contributed less than 2% of relative PEV on average (Fig. 4 supplement 1A).

The high explanatory power of the 14 basis motifs suggests they provide a low-dimensional basis for capturing the dynamics of neural activity in the cortex. This is consistent with the number of motifs (∼19) identified in each recording session (the slightly lower number of basis motifs could reflect spurious noise in individual sessions). Importantly, the number of discovered basis motifs was robust to CNMF parameters (Fig. 4 supplement 1B) and potential hemodynamic contributions to basis motifs were minimal (Fig. 4 supplement 2, see Methods for details). In addition, the low number of basis motifs was not due to the resolution of our approach. We estimated the functional resolution of our imaging approach by correlating pixels across time (Fig. 4 supplement 3). This revealed ∼18 separate functional regions in dorsal cortex (Fig. 4 supplement 3). Individual motifs engaged multiple of these regions over time (Fig. 4C and supplement 3), consistent with the idea that motifs were not constrained by our imaging approach. Indeed, the number of motifs observed was substantially less than the possible number of motifs; even if motifs engaged only 1-2 of these regions, there are still 18^2^=324 different potential motifs, much higher than the 14 we observed. Finally, low dimensionality of basis motifs was not due to compositionality of motifs across time, as this was penalized in the discovery algorithm and the temporal dependency between motifs was weak (Fig. 4 supplement 4; see Methods for details).

### Basis motifs generalize across behaviors

So far, we have only described the motifs of neural activity in animals ‘at rest’. To test whether these same motifs can explain neural activity in multiple behavioral states, we imaged dorsal cortex while animals were engaged in a variety of behaviors.

To begin, we measured the expression of basis motifs in different spontaneous behavioral states. Two mice, which were not used to define the original basis motifs, were imaged for 1 hour while head-fixed on a transparent treadmill (Fig. 5A). As with the original mice, basis motifs captured the majority of variance in neural activity in both animals (Mouse 1: 64.42%, Mouse 2: 66.66%). The ability of basis motifs to generalize outside the set of animals in which they were discovered provides further support for the idea that basis motifs capture core, repeated, spatio-temporal dynamics in neural activity.

**Figure 5.**
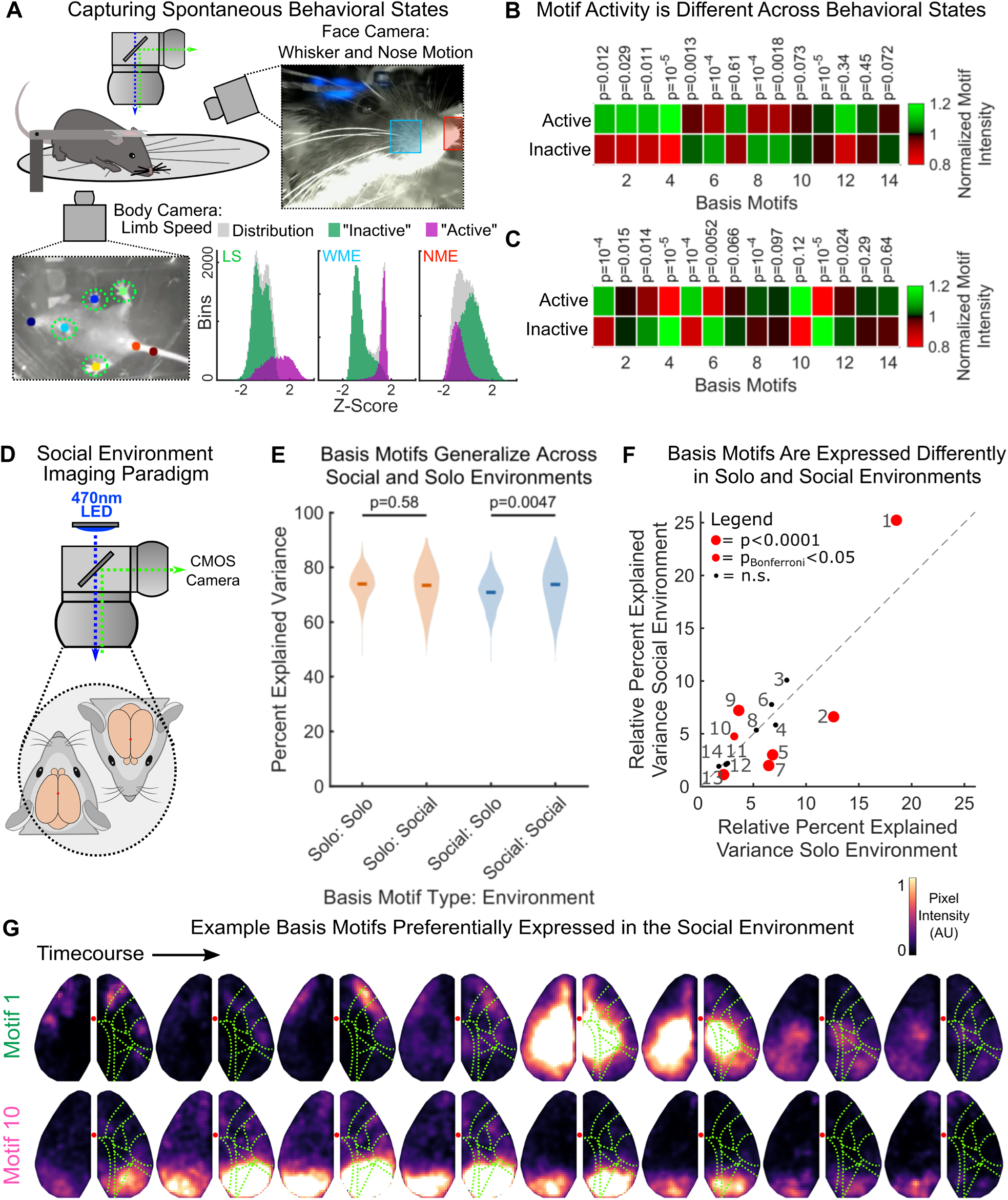
Basis motifs generalize to new animals, across behavioral states, and to social environments. **(A)** Schematic of simultaneous mesoscale imaging and video capture of spontaneous behavior of head-fixed mice on a transparent treadmill. Two new mice, not from the original cohort were used. Blue and red squares indicate area used to measure whisker pad motion energy (WME) and nose motion energy (NME), respectively. Colored dots indicate tracked position of the forelimbs, nose, and tail. These were used to estimate limb speed (LS). Distribution of behavioral variables are shown in three histograms along bottom. Gaussian mixture models fit to the distributions of LS, WME, and NME simultaneously. Two states were discovered: an “active” and “inactive” state (inset; purple and green respectively, see Methods for details). **(B-C)** Motif activity evaluated across different behavioral states in mouse 1 (**B**) and mouse 2 (**C**). Heatmaps show the mean activity of each motif (x-axis) during a behavioral state (y-axis) normalized to the mean activity for that motif across all states. Motif activity was estimated as the mean pixel value of a motif reconstruction over time (see Methods for details). P-values compare motif activity across the two behavioral states. Note: behavioral states across mice are not directly comparable since “active” and “inactive” states reflect a diversity of possible behaviors, as confirmed by manual inspection of videos. **(D)** Schematic of social environment imaging paradigm. **(E)** Comparison of the percent of variance in neural activity that could be explained when animal was alone (at rest, ‘solo’) or when paired with another animal (‘social’). Basis motifs were estimated in each setting and then fit to withheld data in both settings (as in Figure 4C; N=144 and N=123 withheld epochs for solo and social settings, respectively). Labels indicate identification environment: fitting environment (e.g. ‘Solo: Social’ indicates basis motifs from solo environment fit to withheld social data). Full distribution shown; dark lines indicate median. **(F)**. Scatter plot of the relative PEV for each basis motif in the solo environment (x-axis; N=144 epochs) versus the social environment (y-axis; N=123 epochs). Motif labels are indicated with numbers, red markers indicate significant differences in expression rate between environments. Identity line shown along diagonal. **(G)** Example basis motifs preferentially expressed in the social environment. Display and motif labels follow Figure 4. All p-values estimated with Mann-Whitney U-test.

Next, we were interested in whether basis motifs were specific to different behavioral states in the freely behaving mice. Building from recent work, we used infrared cameras to track three different behavior measures: limb speed, whisker pad motion energy, and nose motion energy (Mathis et al., 2018; Musall et al., 2019). To classify behavioral state at each point in time, we fit a gaussian mixed model to the distribution of these three measures (Fig. 5B; see Methods for details). Behavior fell into two broad categories: an ‘active’ state (high whisker pad energy, low nose motion energy, high limb speed) and an ‘inactive’ state (low whisker pad energy, high nose motion energy, low limb speed; Fig 5A). Behavioral states typically lasted for several seconds (median duration for active states: 2.65s and 2.46s; inactive states: 7.88s and 11.54s in mouse 1 and 2 respectively), a longer timescale than the motifs (which are all less than 1 second).

All motifs occurred in both behavioral states (Fig. 5B-C). Follow-up, detailed analyses did not find specific motifs were associated with any specific behaviors, at least as captured by our three tracked measures of behavior (i.e. no motif was associated with grooming, onset of walking, stopping walking, paw repositioning, sniffing, whisking, etc.; data not shown). However, the activity of motifs did vary across the two broad behavioral states (Fig. 5B-C; the activity of 9/14 and 10/14 motifs were different across behavioral states at p<0.05 in Mouse 1 and 2, respectively, Mann-Whitney U-test; this is significantly more than chance, p=10^-11^ and p=10^-12^, binomial test). Similarly, the distribution of motif responses could be used to classify which behavioral state the animal was in (mean classification AUC was 63.42% and 66.51% on withheld data, 99/100 cross-validations were above chance for both animals, 50%; see Methods for details). Therefore, while motifs were not specific to an individual behavioral state, how often a motif was expressed differed across states.

Similar results were seen when animals were engaged in social behaviors. Using a novel paired-imaging paradigm, two mice were simultaneously imaged under the same widefield macroscope (Fig. 5D; see Methods for details). Mice were head-fixed near one another (∼5mm snout-to-snout), enabling sharing of social cues (e.g. whisking, sight, vocalizations, olfaction). To add richness to the sensory environment, mice were intermittently exposed to ‘auditory movies’ that consisted of male mouse vocalizations and synthetic tones (see Methods for details). In this way, the social environment provided a rich, unstructured sensory environment that is fundamentally different from the solo, low-sensory environment used to define the basis motifs. Even in this vastly different environment, basis motifs defined in the original environment captured 73.41% (CI: 71.85-75.23%) of the variance in neural activity (Fig. 5E, right orange plot). This was similar to the variance explained in the solo environment (Fig. 5E, left orange plot; 73.91% CI: 73.14-75.19%; difference between the solo and social environments=0.50%, p=0.49, N=144 solo epochs, N=123 social epochs; Mann-Whitney U-test). Similarly, the opposite was generally true: 11 basis motifs were identified in the social recordings, which captured slightly less of the neural activity in the solo environment (Fig. 5E blue plots; solo=70.83% CI: 69.77-72.09%, social=73.72% CI: 71.54-75.05%; difference=2.89%, p=0.0047, Mann-Whitney U-test).

As before, many basis motifs changed their expression when the behavioral state of the animal changed. Half of the basis motifs had a significant change in their relative explained variance in the social environment compared to baseline (Fig. 5F; 7/14 were different at p_Bonferroni_<0.05, Mann-Whitney U-test; significantly more than chance, p=10^-16^, binomial test). Given the nature of social interactions in mice, one would expect tactile- and visual-associated motifs to be increased. Consistent with this prediction, the motifs elevated in the social environment (motifs 1, 9, and 10) involved somatosensory and/or visual regions (Fig. 5G and 4C).

### Specific motifs capture visual and tactile sensory processing

To begin to understand the computational role of specific motifs, we measured the response of the motifs to tactile and visual stimuli (Fig. 6A, see Methods for details). Solo, awake mice (N=9) were imaged while presented with either moving gratings on a screen or airpuffs to their whiskers (see Methods for details). As before, the distribution of motifs differed between both sensory environments and the original, resting environment; a large proportion of motifs showed a significant change in their relative PEV (Fig. 6B-C: visual: 9/14, tactile: 11/14, significantly different at p_Bonferroni_<0.05, Mann-Whitney U-test; N=1109 visual and N=1110 tactile presentations, N=144 original epochs). Most of these changes were reductions in explained variance, with only a few motifs increasing their expression to the stimuli (Fig. 6B-C).

**Figure 6.**
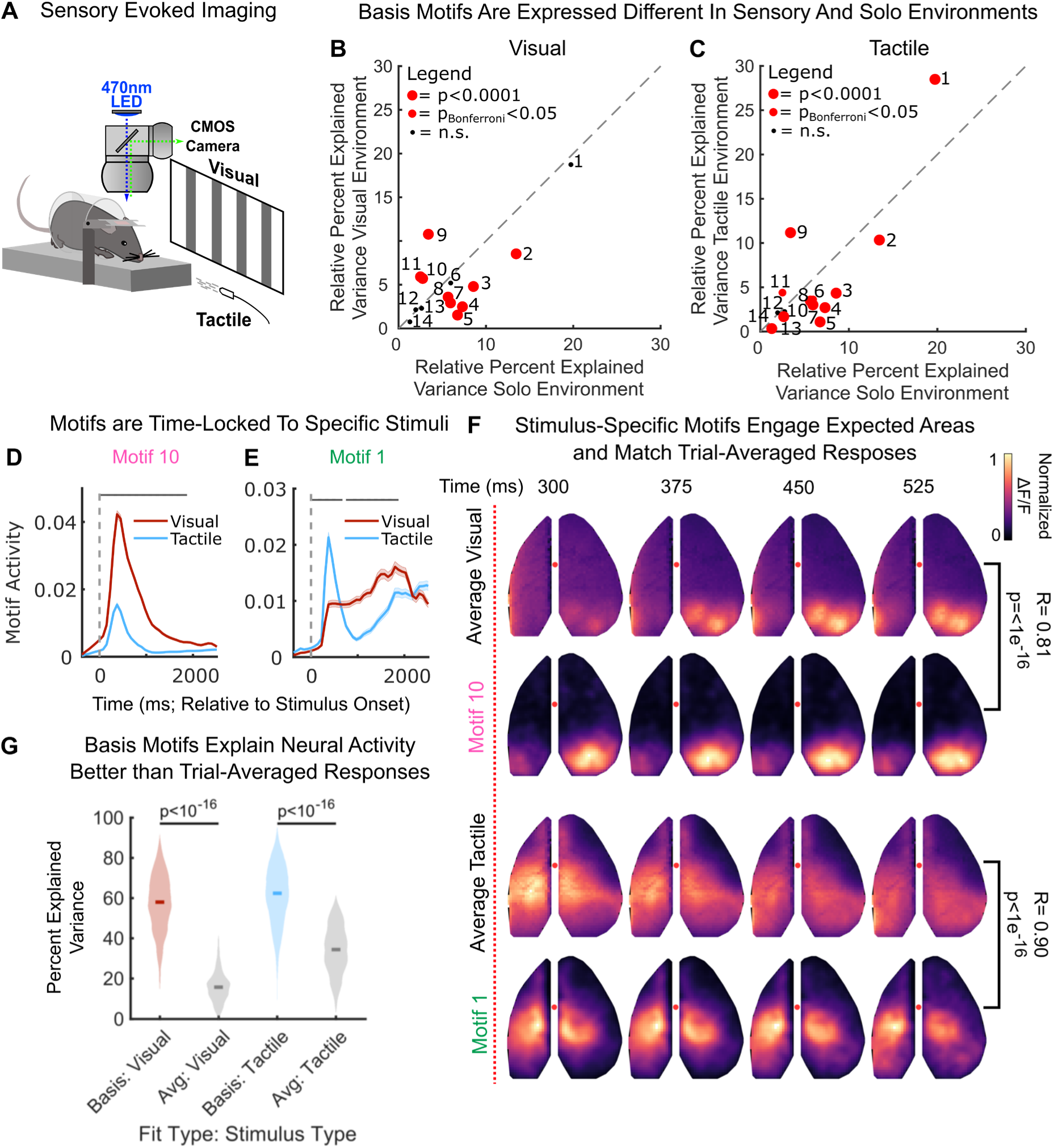
Specific basis motifs reflect processing of specific stimulus modalities. **(A)** Schematic of sensory stimulation paradigm. All stimuli were delivered on animals’ left side. **(B-C)** Scatter plot of the relative PEV for each basis motif in the solo environment (x-axis; N=144) versus the **(B)** visual or **(C)** tactile environment (y-axis; N=1109 and N=1110 visual and tactile samples, respectively). Display follows Figure 5F, showing activation in solo environment at rest (x-axis) versus activation in response to sensory stimuli. Significance computed with Mann-Whitney U-test. **(D-E)** Timecourse of **(D)** motif 10 and **(E)** motif 1 activity relative to stimulus onset (vertical dotted line). Lines and shaded regions indicate mean +/- SEM motif activity in response to visual (red; N=1109) and tactile (blue; N=1110) stimulation. Motif activity calculated as the mean pixel value of a motif reconstruction over time (see Methods for details). Horizontal grey bar indicates significant difference in motif activity between visual and tactile stimuli (p_Bonferroni_ < 0.05, two-sample t-test; see Methods for details). **(F)** Comparison of the trial-averaged stimulus-evoked response and the stimulus-evoked response of the selective basis motifs in **B-C**. First two rows show responses for visual stimuli; third and fourth rows for tactile stimuli. Correlation between average response and basis motif is indicated along right side (pixelwise correlation, Pearson’s ρ). As amplitude of response is arbitrary, pixel intensities were normalized from 0-to-1 before correlation. **(G)** Basis motifs explain more of the variance in neural activity than the average stimulus response. Distributions show the percent of variance in neural activity during the 5 s after stimulus explained by basis motifs (colors) or average responses (gray). Full distribution shown; dark lines indicate median. Significance computed with Wilcoxon Signed-Rank test.

Specific motifs were evoked in response to visual or tactile stimulation. Motif 10 was selectively induced by visual stimulation (Fig. 6D), while motif 1 was selectively induced by tactile stimulation (Fig. 6E). Consistent with a role in sensory processing, both motifs had increased expression in their respective environments (Fig. 6B-C). Furthermore, the spatio-temporal pattern of activity in each motif matched the trial-averaged response to the associated stimulus; motif 10 captured activity in visual cortex, while motif 1 captured activity in motor, somatosensory and parietal cortices (Fig. 6F). Reflecting their overlap, both motifs were significantly correlated with the trial-averaged response (Fig. 6F, and Pearson’s R=0.81, p<10^-16^, for correlation between the average visual response and motif 10, and R=0.90, p<10^-16^, for correlation between the average tactile response and motif 1, all taken during the 13 timepoints post stimulation onset).

While certain basis motifs captured the sensory-evoked neural activity, other motifs were able to capture trial-by-trial ‘noise’ in neural activity. For both stimuli, basis motifs, identified at rest, captured the majority of the variance in neural activity (Fig. 6G; visual: 58.13% CI: 57.11-59.18%; tactile: 62.37% CI: 61.42-63.62%, N=1110 tactile and 1109 visual stimulus presentations; see Methods for details). This was significantly more variance than could be explained by the mean response to each sensory stimulus alone (Fig. 6G; Mean response fits: visual: 17.25% CI: 16.61-17.72%; tactile: 34.42% CI: 33.34-35.40%; difference between motif and mean fits: visual: 40.88%, p<10^-16^; tactile: 27.95%, p<10^-16^; Wilcoxon Signed-Rank Test). This highlights the high variability in responses to a sensory stimulus across trials. Typically, such variability would be discarded as ‘noise’ unrelated to sensory processing. Instead, our results suggest this variability has structure: it is due to the engagement of other motifs that are, presumably, related to other, ongoing, computations.

Finally, we sought to determine whether motifs reflect general stimulus processing or specific stimulus features. To this end, we compared motif expression in response to two visual stimuli (Fig. 7; gratings moving medial to lateral or lateral to medial; see Methods for details and Fig. 7 supplement 1 for similar analysis of tactile stimuli). Unlike comparisons across sensory modalities, the relative percent explained variance in neural activity captured by each basis motif was the same for both visual stimuli (p>0.24 for all 14 motifs; Mann-Whitney U-Test; N=554 and 555 stimulus presentations for medial to lateral and lateral to medial grating respectively). For example, the visually responsive motif 10 responded equally to both stimuli (Fig. 7A). This was not due to limits in spatial resolution of our imaging approach or analytical smoothing. Figure 7B shows pixel-wise classification of the same data can decode stimulus identify (p=0.022, N=9 animals, one-sample t-test; see Methods for details). However, the same classification analysis on data reconstructed from motif activity failed to distinguish between stimuli (p=0.87, N=9 animals, one-sample t-test; difference between classification on original and reconstructed data was significant, p=0.016, paired sample t-test). Thus, the specifics of visual stimuli were encoded in the residuals after fitting the motifs. However, these details contributed minimally to the overall neural activity during the stimulus. The stimulus-specific residuals captured only 3.85% +/- 0.70% SEM of the explainable variance (Fig. 7C). In contrast, motifs captured the vast majority of explainable variance (19.23% +/- 3.23% SEM for stimulus-specific motif 10; 76.92% +/- 3.86% SEM, for remaining motifs; Fig. 7C). Taken together, our results show that motifs capture large-scale patterns of neural activity but are generally agnostic to the finer-grain local activity that captured specifics of stimuli. This is consistent with the idea that motifs capture the broader flow of information across cortical regions.

**Figure 7.**
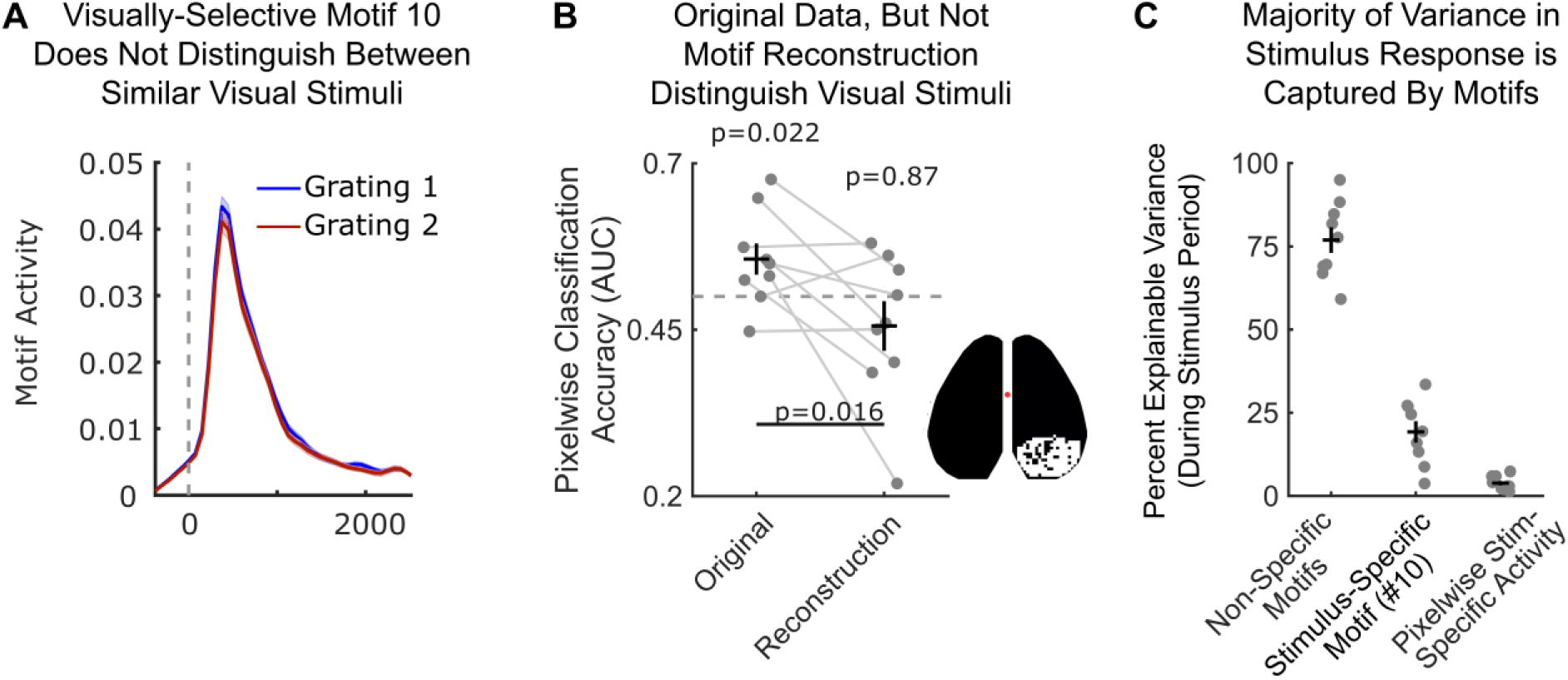
Basis motif 10 reflects general visual stimulus processing. **(A)** Timecourse of motif 10 intensity relative to onset (vertical dotted line) of visual grating 1 (blue; N=554) or visual grating 2 (red; N=555). Display follows Figure 6D-E. No significant differences were observed between stimuli at any timepoint. p>0.11 for all timepoints; two-sample t-test. **(B)** Visual stimuli can be decoded from neural response but not motif response. Classification was done using a support vector machine (SVM) classifier (see Methods for details) and accuracy is shown for withheld validation trials. Markers indicate classifier performance (measured with AUC) for each animal (N=9). Classifiers were trained on either raw pixel values (left column) or reconstructed motif response (right column). Inset shows pixels used for classification (see Methods for details). Dotted line denotes chance (AUC=0.5). Significance computed with one-sample t-test. **(C)** Percent explainable variance in neural activity in response to visual stimuli captured by motifs and stimulus-specific residuals. Stimulus specific residuals are the trial-averaged residuals of motif reconstructions to each visual stimulus type. Data points correspond to mice (N=9). Black bars denote mean (horizontal) and SEM (vertical).

## Discussion

### Spatio-temporal dynamics of cortex-wide activity

Our results show that neural activity is highly dynamic, evolving in both time and space. Leveraging mesoscale calcium imaging in mice, we tracked the spatio-temporal dynamics of neural activity across the dorsal surface of the cortex. Using a convolutional factoring analysis, we identified ‘motifs’ in neural activity. Each motif reflected a different spatio-temporal pattern of activity, with many motifs capturing the sequential activation of multiple, functionally diverse, cortical regions (Figs. 1-3). Together, these motifs explained the large majority of variance in neural activity across different animals (Fig. 4) and in novel behavioral situations (Fig. 5-6).

A couple of the motifs captured patterns of activity observed in previous work, supporting the validity of the CNMF approach. For example, previous work has studied spatio-temporal waves of activity that propagate anterior-to-posteriorly across the cortex at different temporal scales (Greenberg et al., 2018; Matsui et al., 2016; Mitra et al., 2018). Our motifs (1 and 11) recapitulate these waves, along with their temporal diversity (motif 1 = fast, motif 11 = slow). In addition to these motifs, we also discovered several additional, spatio-temporally distinct, anterior-to-posterior propagating waves (motifs 2, 4, and 9).

Similarly, brain regions that were often co-activated in motifs were aligned with previously reported spatial patterns of co-activation in the mouse cortex (Musall et al., 2019; Silasi et al., 2016; Vanni et al., 2017). For example, motifs 6, 8, 11, and 14 include coactivation of anterolateral somatosensory and motor regions. This pattern is observed often and reflects the close functional relationship between motor activity and somatosensory processing. Here, we extend this work by showing neural activity can flow within and between these networks in different ways.

Relatedly, previous work using mesoscale imaging demonstrated that the mouse cortex exhibits repeating patterns of activity (Mohajerani et al., 2013). However, this work relied on identifying average patterns evoked by sensory stimuli (visual, tactile, auditory) and correlating the spatially and temporally static templates of those patterns to activity in resting animals. As we demonstrate, stimulus-evoked patterns capture considerably less variance in neural activity than our approach, even in response to sensory stimuli themselves (average stimuli responses: ∼15-35% of activity versus motifs: ∼60%). In addition, previous work has used zero-lag correlations to show the brain transitions through different functional network states over time (Ashourvan et al., 2017; Hutchison et al., 2013; Preti et al., 2017; Vidaurre et al., 2017). Here, we show that these functional network states themselves have rich dynamics, reflecting specific sequential patterns of activity across the network. By encapsulating these dynamics, motifs are able to capture significantly more of the variance in neural activity compared to static networks.

### Motifs of neural activity may reflect cognitive computations

Each motif captured a different spatio-temporal pattern of neural activity. As neural activity passes through the neural network of a brain region, it is thought to be functionally transformed in a behaviorally-relevant manner (e.g. visual processing in visual cortex or decision-making in parietal cortex). Therefore, the dynamic activation of multiple regions in a motif, could reflect a specific, multi-step, ordered transformation of information. In this way, the basis motifs would reflect a set of ‘core computations’ carried out by the brain.

Consistent with this hypothesis, the distribution of motifs differed across behavioral states (Fig. 5) and in response to social and sensory stimuli (Figs. 6-7). Specific motifs were also associated with specific cognitive processes, such as tactile and visual processing (Fig. 7). This follows previous work showing the engagement of brain networks is specific to the current behavior (Mattar et al., 2015; Telesford et al., 2016) and that disrupting these networks underlies numerous pathologies (Badhwar et al., 2017; Braun et al., 2016; Harlalka et al., 2019).

In this way, our results are consistent with a hierarchical relationship between behavior and cortical dynamics. Behavior engages a set of core cognitive computations, reflected in the activation of multiple motifs in each behavioral state (Fig. 5). Multiple behavioral states engage the same motifs, but they engage them to a different extent. This suggests different behavioral states do not require independent patterns of neural activities, but rather reflect a re-weighting of the expression of core computations.

### Low dimensional structure of cortex-wide dynamics may facilitate cognitive control of neural activity

Our results show that the dynamics of the cortex are low dimensional. Motifs identified in different animals and recording sessions clustered into a limited set of 14 unique ‘basis’ motifs. This limited number of basis motifs captured the large majority of variance in neural activity (∼75%) across animals, across behavioral states, and in different behavioral environments. Such a low-dimensional repertoire of cortical activity is consistent with previous work using zero-lag correlations of neural activity to measure functional connectivity between brain regions (e.g. 17 functional networks in the human cortex using fMRI, Yeo et al., 2011).

Why might cortex-wide neural activity be low dimensional? Given the multitude of sensory inputs, internal states, and motor actions, one might expect neural activity to be extremely high dimensional. Indeed, recent large-scale electrophysiology and imaging studies have found high dimensional (>100) representations within a brain region, with neural activity often representing small aspects of behavior (e.g. facial twitches, limb position, etc; Stringer et al., 2019a, 2019b; Lieber and Bensmaia, 2019). However, while this high dimensionality of neural representations is great for capturing the fullness of an experience, its complexity presents a problem for cognitive control.

In most situations, we do not *a priori* know the exact set of computations to engage. Instead, we must learn which computations are optimal for the current situation. One way to do this is to search through the space of possible computations until the ‘best’ computation is found. This could explain the diversity of motifs observed in our animals at rest; the expression of different motifs reflects the ‘sampling’ of different computations as the animals search for the one that fits their current, novel, situation. Similarly, the greater consistency in the motifs expressed during social or sensory environments may reflect a reduction in the uncertainty about what computations to engage in.

However, in this framework, a high dimensionality is costly. As the number of possible computations (and their neural representations) increases, it becomes harder to find the optimum (the classic ‘curse of dimensionality’ problem; Bellman, 2003). Therefore, limiting the number of motifs may make it easier to find a contextually appropriate computation. The caveat to a low dimensional repertoire of motifs is that it necessitates a coarser sampling of computational space, which may limit how well the best computation approximates the true optimal computation. In this way, the dimensionality of dynamics may reflect a trade-off between the time it takes to identify an appropriate computation and the optimality of that computation.

More broadly, our results are consistent with a hierarchical control of information processing in the brain (Mearns et al., 2019; Deco and Kringelbach, 2017; Ashourvan et al., 2019; Park and Friston, 2013; Botvinick, 2008). Behavioral state changes slowly and reflects the animal’s broad behavioral goal (Wiltschko et al., 2015). On a shorter timescale, control mechanisms coordinate broad dynamics across cortical regions (motifs). This allows the animal to engage in a behaviorally relevant category of computations (e.g. tactile versus visual processing). High dimensional representations in local circuits then produce more nuanced processing (e.g. the specific identity of a visual stimulus). In this way, by activating specific motifs, control processes could direct the broader flow of information across brain regions (a low-dimensional problem) while avoiding the difficulty of directing detailed single-neuron processing (a high-dimensional problem).

### Future Directions

Our approach has several limitations that motivate future research. First, although we are able to capture a large fraction of the cortex, we are limited to the dorsal cortex and so miss out on lateral auditory cortex, cortical regions deep along the midline, and all sub-cortical regions. Second, while we probed activity in several different environments, imaging was always restricted to head-fixed animals. Third, the number and identity of motifs, as well as the relative contributions of spatial and temporal dynamics to variance in neural activity, is likely influenced by the nature of our approach. The relatively slow timecourse of GCaMP6f (Chen et al., 2013), and biases in the neural activity underlying the calcium signal (Allen et al., 2017; Makino et al., 2017) may have decreased the spatial resolution and slowed the temporal dynamics. However, it is important to note that the spatial resolution used in our imaging approach (∼136 µm^2^/pixel) is higher than the broad activation of brain regions observed in the motifs and was high enough to capture pixel-specific information about stimuli beyond the motifs. Furthermore, our functional resolution was able to distinguish 18 distinct regions. Motifs engaged multiple regions, suggesting motifs were broader than the functional resolution (and therefore not limited by our approach). Finally, even given these constraints, the number of potential spatio-temporal patterns is far higher than the 14 basis motifs found.

In addition to addressing these limitations, future work is needed to understand the computations associated with each motif. Here we’ve used simple behavioral paradigms to associate a few motifs with sensory processing. However, the computation underlying many of the motifs remains unknown – by cataloging motif expression across experiments and behaviors, we can begin to understand the function of each motif and gain a more holistic understanding of how and why neural activity evolves across the brain in support of behavior.

## Acknowledgments

The authors thank Lucas Pinto, Morgan Gustison, Marcelo Mattar, Chantal Stern, Alex Libby, Matt Panichello, Sina Tafazoli, Caroline Jahn, Flora Bouchacourt, Emily Dennis, and Sarah Henrickson for their detailed feedback during the writing of this manuscript. We also thank Stephan Thiberge for designing and constructing the widefield macroscope. We thank the Princeton Laboratory Animal Resources staff for their support. This work was funded by NIH DP2 EY025446.

## Material and Methods

### Key Resources Table

| REAGENT or RESOURCE | SOURCE | IDENTIFIER |
| --- | --- | --- |
| Experimental Models: Organisms/Strains |  |  |
| Mouse: Thy1-GCaMP6f:<br>*C57BL/6J-Tg(Thy1-GCaMP6f)GP5.3Dkim/J* | The Jackson Laboratory | #028280 |
| Software and Algorithms |  |  |
| MATLAB 2017a-2018b | Mathworks | N/A |
| seqNMF MATLAB Toolbox | Mackevicius et al., 2019 | N/A |
| MUPET MATLAB Toolbox | Van Segbroeck et al., 2017 | N/A |
| OFAMM MATLAB Toolbox | Afrashteh et al., 2017 | N/A |
| Psychtoolbox-3 MATLAB Toolbox | http://psychtoolbox.org/<br>(Brainard, 1997) | N/A |
| Python version 3.6.4 | Python Software Foundation | N/A |
| DeepLabCut Python Library | Mathis et al., 2018 | N/A |
| Signal Processing and Analysis code for<br>Widefield Imaging data | This paper | N/A |

### Lead Contact and Material Availability

Further information and request for resources and reagents should be addressed to Lead Contact, Timothy J. Buschman

### Data and Software Availability

Preprocessed data is available on Dryad data repository as image stacks (saved in Matlab file format; DOI: 10.5061/dryad.kkwh70s1v; url: https://datadryad.org/stash/share/-q39l6jEbN--voeSe2-5Y3Z1pfbeYEFLO-Kf_f-cjpE). The data has been preprocessed as described below (spatially binned, masked, filtered, and then thresholded). Due to file size constraints, the full raw data is not available on the Dryad repository but is available upon request. Example data and figure generation code will be available on GitHub (https://github.com/buschman-lab) upon acceptance.

### Experimental Model

All experiments and procedures were carried out in accordance with the standards of the Animal Care and Use Committee (IACUC) of Princeton University and the National Institutes of Health. All mice were ∼6-8 weeks of age at the start of experiments. Mice (N=11) were group housed prior to surgery and single housed post-surgery on a reverse 12-hr light cycle. All experiments were performed during the dark period, typically between 12:00 and 18:00. Animals received standard rodent diets and water *ad libitum*. Both female (N=5) and male (N=6) mice were used. All mice were C57BL/6J-Tg(Thy1-GCaMP6f)GP5.3Dkim/J (The Jackson Laboratory; Dana et al., 2014). 9 mice were used for solo (rest) and sensory environment widefield imaging experiments. These mice were control animals from a larger study. In that context, these animals were the offspring of female mice that received a single intraperitoneal injection (0.6-0.66mL, depending on animal weight) of sterile saline while pregnant. A subset of these mice (N=7) were used for social imaging experiments. Separate mice (N=2) were used for spontaneous behavioral state and hemodynamic correction experiments (these mice did not receive *in utero* exposure to saline).

### Surgical Procedures

Surgical procedures closely followed Guo et al. (2014). Mice were anesthetized with isoflurane (induction ∼2.5%; maintenance ∼1%). Buprenorphine (0.1mg/kg), Meloxicam (1mg/kg), and sterile saline (0.01mL/g) were administered at the start of surgery. Anesthesia depth was confirmed by toe pinch. Hair was removed from the dorsal scalp (Wahl, Series 8655 Hair Trimmer), the area was disinfected with 3 repeat applications of betadine and 70% isopropanol, and the skin removed. Periosteum was removed and the skull was dried. A thin, even layer of clear dental acrylic was applied to the exposed bone and let dry for ∼15 minutes (C&B Metabond Quick Cement System). Acrylic was polished until even and translucent using a rotary tool (Dremel, Series 7700) with rubber acrylic polishing tip (Shofu, part #0321). A custom titanium headplate with a 11mm trapezoidal window was cemented to the skull with dental acrylic (C&B Metabond). After the cement was fixed (∼15 minutes), a thin layer of clear nail polish (Electron Microscopy Sciences, part #72180) was applied to the translucent skull window and allowed to dry (∼10 minutes). A custom acrylic cover screwed to the headplate protected the translucent skull after surgery and between imaging sessions. After surgery, mice were placed in a clean home cage to recover. Mice were administered Meloxicam (1mg/kg) 24 hours post-surgery and single housed for the duration of the study.

### Widefield Imaging

Imaging took place in a quiet, dark, dedicated imaging room. For all experiments except spontaneous behavioral monitoring (detailed below), mice were head-fixed in a 1.5 inch diameter × 4 inch long polycarbonate tube (Fig. 1A) and placed under a custom-built fluorescence macroscope consisting of back-to-back 50 mm objective lens (Leica, 0.63x and 1x magnification), separated by a 495nm dichroic mirror (Semrock Inc, FF495-Di03-50×70). Excitation light (470nm, 0.4mW/mm^2^) was delivered through the objective lens from an LED (Luxeon, 470nm Rebel LED, part #SP-03-B4) with a 470/22 clean-up bandpass filter (Semrock, FF01-470/22-25). Fluorescence was captured in 75ms exposures (FPS = 13.3Hz) by an Optimos CMOS Camera (Photometrics). Prior to imaging, the macroscope was focused ∼500um below the dorsal cranium, below surface blood vessels. Fluorescence activity was captured at 980×540 resolution (∼34um/pixel) when the animal was imaged alone.

Images were captured using Micro-Manager software (version 1.4, Edelstein et al., 2014) on a dedicated imaging computer (Microsoft, Windows 7). Image capture was triggered by an analog voltage signal from a separate timing acquisition computer. Custom MATLAB (Mathworks) code controlled stimulus delivery, recorded gross animal movement via a piezo sensor (SparkFun, part #09197) attached to the animal holding tube, and captured camera exposure timing through a DAQ card (National Instruments, PCIe-6323 X Series, part #7481045-01). Timing of all camera exposures, triggers, behavioral measures, and stimulus delivery were captured for post-hoc timing validation. No frames were dropped across any imaging experiments. A camera allowed remote animal monitoring for signs of distress.

For recordings of spontaneous cortical activity, mice were head-fixed in the imaging rig and habituated for 5 minutes. After habituation, cortical activity was recorded for 12 consecutive minutes and stored as 3, 4-minute stacks of TIFF images. Qualitative real-time assessment of behavioral videos and post-hoc analysis of activity (captured by piezo sensor) revealed minimal episodes of extensive motor activity (e.g. struggling) during imaging. As our goal is to capture all behavioral states, we did not exclude these moments from our analysis. Instead, motifs captured these events alongside other cortical events.

### Widefield Imaging: Spontaneous Behavioral State Monitoring

Widefield imaging was performed as above with minor modifications. Mice were head-fixed on a custom transparent acrylic treadmill and illuminated with infrared light (Univivi 850nm IR Illuminator). During imaging, behavioral measures were captured using two cameras: a PS3 EYE webcam (640×480 pixel resolution) focused on the animal’s whole body and a GearHead webcam (320×240 pixel resolution) focused on the animal’s face. Custom python (v3.6.4) scripts synchronized the frame exposure of the behavioral cameras at 60Hz.

### Widefield Imaging: Paired Social Environment

The macroscope objectives from the above widefield imaging paradigm were replaced with 0.63x and 1.6x magnification back-to-back objectives, permitting an ∼30×20mm field of view (lens order: mouse, 0.63x, 1.6x, CMOS camera). Images were acquired at 1960 × 1080 resolution (∼34um/pixel). Animals were precisely positioned to be the same distance from the objective. Mice faced one another, approximately eye-to-eye in the anterior-posterior axis. Their snouts were separated along the medial-lateral axis by a 5-7 mm gap; close enough to permit whisking and social contact but prevent adverse physical interactions. A 1mm plexiglass divider at snout level ensured no paw/limb contact. Mice were positioned in individual plexiglass tubes. Pairs were imaged together for 12 consecutive minutes, once each recording day. Some pairings included mice outside this study cohort. 76 recordings from the experimental cohort were collected. After each recording, the imaging apparatus was thoroughly cleaned with ethanol and dried before imaging of the next pair (removing olfactory cues).

Animal pairs were provided with sensory stimuli consisting of playback of pre-recorded, naturalistic ultrasonic vocalizations (USVs) between adult mice, synthetic USVs, or ‘background’ noise. Naturalistic USV stimuli were obtained from the mouseTube database (Torquet et al., 2016). In particular, we used four recordings of 3 min interactions between male and estrus female wildtype C57BL/6J mice (files S2-4-4, S2-4-105, S2-4-123, S2-4-138). Details on the methods used to record these interactions are described in the original study by Schmeisser et al. (2012). To produce more salient stimuli, we reduced these 3 min recordings into 1 min recordings by using Praat software (version 6.0.23) to shorten the silent periods between USV bouts. We bandpass filtered these recordings to the 40-100 kHz range (Hann filter with 100 Hz smoothing) to reduce extraneous background noise, and down sampled the recordings to 200 kHz.

Synthetic USV stimuli were generated using a customized MATLAB script that created artificial sine wave tones matching the spectro-temporal properties of naturalistic stimuli. Specifically, synthetic stimuli had the same rate (calls per minute), average duration, and mean frequency as naturalistic USVs (we used MUPET to characterize USV properties; Van Segbroeck et al., 2017). Tones were evenly spaced throughout the synthetic stimulus. Background noise was generated from the silent periods of the 3-minute vocalization recordings. Each recording session contained 1 epoch of naturalistic USVs, 1 epoch of synthetic USVs, and 1 epoch with background noise. All acoustic stimuli were presented at ∼70dB through a MF1-S speaker (Tucker Davis Technologies) placed 10cm away from both subjects.

### Widefield imaging: Structured Sensory Environments

Widefield imaging was performed as in the original (solo) condition. All stimuli were provided to the animals’ left side. Recordings were 15-minutes long, divided into 90 trials of 8000ms duration. Trials were structured with a 3000ms baseline period, 2000ms stimulus period, and 3000ms post-stimulus period. Trials were separated by an inter-trial interval randomly drawn between 1000 and 1750ms.

Air puffs (10psi) were gated by solenoids (NResearch, Solenoid valve, part #161K011) and were directed at the whisker pad in either the anterior-to-posterior or posterior-to-anterior direction. Visual stimuli were gratings of 2.1cm bar width, 100% contrast, delivered on a 10inch monitor (Eyoyo, 10-inch 1920×1200 IPS LED), positioned 14cm away from animals’ left eye. Gratings drifted from medial-to-lateral or lateral-to-medial at 8 cycles per second. Visual stimuli were presented for 2000ms. During these recordings, mice also received trials of auditory stimuli (e.g. 2 tones). These data were not analyzed since auditory cortex was not imaged and so no evoked response was observed. Each recording captured 30 trials of each stimulus modality. 3329 trials were captured in total across 9 animals: resulting in 1110 tactile, 1109 visual (and 1110 auditory) trials. 1 visual trial was lost due to timing issues.

Due to light-artifact of visual stimuli leaking through the ipsilateral cortical bone in a subset of recordings, a more conservative mask on the ipsilateral hemisphere was used for all sensory environment analyses (as shown in Fig. 6F). Accordingly, this mask was used for all analyses in Figures 6-7, including quantification of motifs in the original (solo) environment.

### Statistical Analysis

All analyses were performed in MATLAB (Mathworks). Number of mice used was based on previously published studies (Makino et al., 2017; Allen et al., 2017; Mohajerani et al., 2013). As described throughout methods and main text, analyses were performed on 11 separate animals, across multiple recording sessions, and 4 behavioral environments (e.g. biological replicates). Analyses were validated across a range of processing parameters (e.g. technical replicates). All statistical tests, significance values, and associated statistics are denoted in the main text. P-values below machine precision are reported as p<10^-16^. All 95% Confidence intervals were computed using MATLAB *bootci* function (1000 bootstrap samples).

### Widefield Imaging Preprocessing

Image stacks were cropped to a 540×540 pixel outline of the cortical window. Images were aligned within and across recordings using user-drawn fiducials denoting the sagittal sinus midline and bregma for each recording. For anatomical reference (Figs. 1A and supplement 1), recordings were aligned to a 2D projection of the Allen Brain Atlas, version CCFv3 using bregma coordinates (Oh et al., 2014; ABA API interfacing with MATLAB adapted from https://github.com/Sainsbury *WellcomeCentre/AllenBrainAPI*). The complete 2D projection is shown in Fig. 1 Supplement 1. As they are only intended to be local references, the parcel outlines overlaid in Figures 1, 4, 5 were created by manually tracing this 2D projection (Inkscape Vector Graphics Software).

Changes in fluorescence due to hemodynamic fluctuations may confound the neural activity captured by widefield imaging (Allen et al., 2017; Ma et al., 2016a, 2016b). However, previous work has found hemodynamic contributions to fluorescent signal using similar widefield imaging approaches are minimal (Cramer et al., 2019; Murphy et al., 2016; Vanni and Murphy, 2014), and can be mitigated by removing pixels corresponding to vasculature (Makino et al., 2017). To mitigate impact of hemodynamic contributions, we masked pixels corresponding to vasculature. To identify vasculature, the middle image of each recording was smoothed with a 2D median filter (neighborhood 125 pixels^2^) and subtracted from the raw image. As vasculature pixels are much darker than pixels of neural tissue, we created a vasculature mask by thresholding the reference image to pixels intensities >= 2.5 standard deviations below the mean. To remove noise, the mask was morphologically closed with a 2-pixel disk structuring element. A vasculature mask was created for each recording. Supplemental experiments, outlined below, demonstrated that vascular masks successfully mitigated the contribution of hemodynamics to our signal.

Vasculature masks were combined with a manually drawn outline of the optically accessible cortical surface and applied to each recording to conservatively mask non-neural pixels. Masks removed the sagittal sinus, mitigating vascular dilation artifacts. Additionally, masks removed peripheral lateral regions, such as dorsal auditory cortex, where fluorescence contributions across animals may be differentially influenced by individual skull curvature. After alignment and registration, recordings were spatially binned to 135×135 pixels (∼68µm^2^/pixel). Masked pixels were ignored for spatial binning. Normalized activity was computed as change in fluorescence, e.g. ΔF/F over time according to 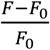. Baseline fluorescence, F_0_ was computed using a 9750ms (130 timepoints) rolling mean. To remove slow fluctuation in signal (e.g. due to change in excitation intensity), pixels traces were detrended using linear least squares fit.

Recordings were bandpass filtered at 0.1 to 4Hz (10^th^ order Butterworth filter). Pixel traces were thresholded at 2 standard deviations per pixel in order to remove noise in spontaneous activity. Values below this threshold were set to zero. Thresholding had minimal impact on subsequent analyses and conclusions. For example, similar numbers of basis motifs (∼14) were discovered in non-thresholded data. After filtering and thresholding, recordings were spatially binned again to a final size of 68×68 pixels (∼136µm^2^/pixel). Pixels with zero variance during an epoch (e.g. masked pixels) were ignored for all subsequent analyses. Subsequent factorizations require non-negative pixel values so recordings where normalized to range of 0 to 1 using the maximum and minimum pixel values per recording. The 12-minute solo and social recordings were divided into six, 2-minute epochs; alternating epochs were used for motif discovery or withheld for testing (Fig. 1B). For solo recordings this resulted in 144 ‘discovery’ and 144 ‘withheld’ epochs.

For social recordings, cortices of individual animals were cropped to 540×540 pixels and preprocessing followed as above. Again, recordings were divided into 2-minute epochs, resulting in a total of 228 ‘discovery’ and ‘withheld’ epochs. Given the proximity of the animals, whiskers from one animal sometimes entered the imaging field of view of the paired animal, creating artifacts easily detected upon manual inspection. All epochs were manually inspected and epochs with any whisker artifacts (N=105) were removed, resulting in 123 ‘discovery’ and ‘withheld’ epochs (8.2 hours in total).

For sensory trials, which were 8 seconds in length, each trial’s ΔF/F was calculated using the mean of the first 26 timepoints (∼2s) as baseline fluorescence. Burst in activity were discovered by thresholding traces at 1 standard deviation per pixel. All other preprocessing steps were followed as above.

### Multiwavelength Hemodynamic Correction

Additional experiments using multiwavelength hemodynamic correction were performed to confirm that vasculature masking mitigated hemodynamic contributions to motifs (Fig. 4 supplement 2). Hemodynamic correction followed (Musall et al., 2019). In brief, widefield imaging was performed while strobing between illumination with a blue LED (470nm, 0.4mW/mm^2^) and violet LED (410nm, LuxDrive LED, part #A008-UV400-65 with a 405/10 clean-up bandpass filter; Edmund Optics part #65-678). Each exposure was 35.5ms and light from both LEDs were collimated and coupled to the same excitation path using a 425nm dichroic (Thorlabs part #DMLP425). Illumination wavelengths alternated each frame. Strobing was controlled using an Arduino Due with custom MOSFET circuits coupled to frame exposure of the macroscope (as in Pinto et al., 2019). After vasculature masking and spatial-binning to 135×135 pixels, violet-exposed frames (e.g. non-calcium dependent GCaMP6f fluorescence, Lerner et al., 2015) were rescaled to match the intensity of blue-exposed frames. ΔF/F was then computed as ΔF/F_blue_ - ΔF/F_violet_. Remaining preprocessing steps followed original (solo) experiments.

### Motif Discovery

We used the *seqNMF* algorithm (MATLAB toolbox from Mackevicius et al., 2019) to discover spatio-temporal sequences in widefield imaging data. This method employs convolutional non-negative matrix factorization (CNMF) with a penalty term to facilitate discovery of repeating sequences. All equations below are reproduced from the main text and Tables 1 and 2 of Mackevicius et al (2019). For interpretability, we maintained the nomenclature of the original paper where possible.

We consider a given image as a *P × 1* vector of pixel values and a recording image sequence (i.e. recording epoch) as a *P × T* matrix, where *T* is the number of timepoints in the recording. This matrix can be factorized into a set of *K* smaller matrices of size *P × L* representing short sequences of events (e.g. *motifs)*. Collectively this set of motifs is termed *W* (a *P × K × L* tensor).

Each pattern is expressed over time according to a *K × T* temporal weighting matrix termed *H.* Thus, the original data matrix can be approximated as the sum of K convolutions between the motifs in *W* and their corresponding temporal weightings in *H*:

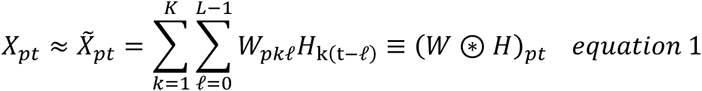

Here, ⊛ indicates the convolution operator. The values of W and H were found iteratively using a multiplicative update algorithm. The number of timepoints for each motif was set to 13 frames (975ms). This *L* value was chosen because it is well above the duration of GCaMP6f event kinetics and qualitative assessment of imaging recordings suggested most spontaneous events were < 1000ms in duration, agreeing with previous literature (Stringer et al., 2019a). Active timepoints in motifs may be shorter than *L*; in which case unused timepoints are zero-padded. Preliminary tests at *L > 13* produced similar results to *L*=13, and explained variance captured by motifs reconstructions plateaued at *L* values >=13 (Fig. 1 supplement 2C). As increasing *L* increased computation time, *L=*13 was used for practical purposes.

The maximum number of possible motifs (*K*) per recording was chosen by iteratively sweeping a range of *K* values and evaluating the number of basis motif clusters at each value (see below regarding basis motif clustering). Figure 4 supplement 1B shows the number of basis motifs asymptotes at *K* >= *12*. K = 28 was used for all motif discovery experiments. *K* = 28 was chosen as it was well above the elbow of the asymptote and produced a total number of basis motifs on the upper range of the test *K* values (thus not analytically limiting the observe dimensionality). As with *L,* if fewer than *K* motifs were found, the remaining motifs would be populated with zeros. Again, we constrained *K* because computational demands when adding extra, unnecessary (blank) motifs.

The *seqNMF* algorithm improves upon typical CNMF by including a spatio-temporal penalty term into the cost function of the multiplicative update algorithm. In brief, this reduces redundancy between motifs: 1) multiple motifs do not describe the same sequence of activity; 2) a single motif is not temporally split into separate motifs; and 3) motifs are encouraged to be non-overlapping in time. This penalty termed is implemented as follows:

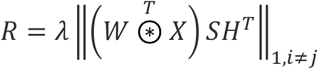

Here, temporal overlap (correlation) between motifs is captured by *SH^T^*. *S* is a *T × T* temporal smoothing matrix where *S_ij_* = 1 when |*i* - *j*| < *L*; otherwise *S_ij_* = 0. Thus, each temporal weighting in *H* is smoothed by a square window of length 2*L*-1, increasing the product of motifs that temporally overlap within that window.

Competition between spatio-temporal structure of motifs is achieved by calculating the overlap of motifs in *W* with the original data as follows

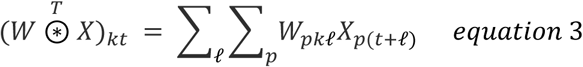

Motifs containing similar patterns will overlap with the original data matrix at the same times. The notation || · ||_1*,i* ± *j*_ ignores penalizing terms along the diagonal such that spatial and temporal autocorrelation of motifs is not penalized. *λ* is a tunable parameter that controls the magnitude of the penalty term. The result of this penalty, when implemented as a cost function in the multiplicative update algorithm for fitting *H* and *W*, is to bias factorization such that only one motif is active at a given timepoint.

We followed the approach used by Mackevicius et al., to determine the magnitude of the *λ* penalty term. *λ* was swept across 6 orders of magnitude from 10^-6^ to 1 (Fig. 1 supplement 2A). The impact on reconstruction cost, motif spatio-temporal correlation, explained variance, and number of factors was evaluated. A value slightly above the cross-over point in reconstruction cost and explained variance was chosen for subsequent experiments (Fig. 1 supplement 2A). Importantly, a similar number of motifs were found for all values of λ, suggesting our results did not depend on the exact value.

Additionally, the *seqNMF* algorithm contains optional orthogonality constraints to bias factorizations towards parts-based and events-based factorizations. In widefield imaging data, parts-based would preferentially detect spatially independent motifs. Events-based factorizations, used in this study, would preferentially discover temporally independent motifs that correspond to specific instantiations of sequential activity patterns. This was achieved with an additional smoothed orthogonality cost term penalizing overlap in motif temporal weightings:

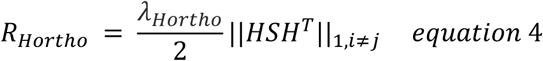

The magnitude of *λ_orthoH_* was set to 1 based on preliminary experiments and remained unchanged for all motif discovery experiments, including the parameter sweeps of *λ* and *K* described above. All fitting processes were run for 300 iterations; at which point the cost function leveled out (Fig. 1 supplement 2B).

Supplemental Table 1 contains a complete list of all adjustable parameter values of the seqNMF algorithm toolbox used for each experiment. Descriptions for parameters not discussed above can be found in original work. To discover motifs in paired social recordings, the same *K* and *L* values (28 and 13) were used. λ was refit following the procedure described above.

Note, one change was made to the *seqNMF* algorithm toolbox. For convenience, the smoothing matrix S was multiplied by 0.01. Specifically, line 98 of original seqNMF code: *smoothkernel = zeros(1,(2*L)-1)* replaced with *smoothkernel = 0.01*zeros(1,(2*L)-1).* This allowed *λ* values to be 100x larger and, therefore, easier to read.

### Visualizing Motif Activity Flow

To aid in visualization of motif patterns (Figs. 1 and 3), we used the Horn-Schunck optical flow method to calculate the velocity vector fields between subsequent timepoints of motifs (Horn and Schunck, 1981; implemented using *HS* function from OFAMM MATLAB toolbox, Afrashteh et al., 2017). For basis motif images, optical flow was only performed for timepoints with non-zero variance across pixels. For all plots, arrows depict the direction and velocity of flow for the top 50% intensity pixels of each timepoint. The number of arrows were downsampled by a factor of three for clarity in visualization.

### Comparing Motifs to sPCA and sNMF

Data preprocessing for Motif, sPCA, and sNMF discovery was identical. For sPCA, pixels were treated as variables and timepoints as observations. sNMF was performed by setting *L* to 1 (effectively reducing *equation 1* to matrix multiplication) and removing all temporal and spatial sparsity terms.

### Refitting Motifs to Withheld Data

To refit motifs to withheld data, we used the same *seqNMF* algorithm, but *W* was fixed to the previously discovered motifs (or basis motifs) during fitting. Thus, the only updatable features of the factorization were motif temporal weightings, *H*. See Supplemental Table 1 for a list of all parameters used during fitting. Importantly, since only temporal weightings were refit, *λ* was set to zero. Thus, unlike motif discovery where motifs were highly spatially and temporally independent, refit motifs did not have this constraint and thus some combinatorial motif activation was observed. However, increasing λ_*Hortho*_ had little impact on the percent of variance explained when refitting the motifs, suggesting that any compositionality had minimal impact on results (Fig. 3 supplement 1). Importantly, all motifs, static networks, and time-varying networks were refit to withheld data in the same way and therefore their percent explained variance can be directly compared.

For solo and social recordings, the part-based factorization orthogonality bias was maintained as in initial discovery (see Supplemental Table 1). Given the short duration of sensory trials, the orthogonality bias was not used. The same process was followed for refitting basis motifs and social basis motifs to withheld data. For all refitting processes, motifs and withheld data were first spatially smoothed with a 2D gaussian filter (σ = [1,1]). Only pixels with non-zero variance in both motifs and withheld data were used.

### Generating Basis Motifs

To identify basis motifs, we used an unsupervised clustering algorithm (Phenograph, Nicosia et al., 2009; Levine et al., 2015). For clustering, motifs were renormalized to 0-to-1 and spatiotemporally smoothed with a 3D gaussian filter (σ = [1,1,0.1]). The Phenograph algorithm generates a directed graph, where each node is connected to its *k* nearest neighbors. Louvain community detection is then performed on this graph to cluster nodes into groups. For finding neighbors, distances between motifs were computed as the peak in their temporal cross-correlation. The only tunable parameter of Phenograph is the number of nearest neighbors (*k*)*. k*=15 was chosen based on initial experiments. Similar number of clusters (10-20) were observed for *k*=10 and *k*=20.

Basis motifs were generated by taking the mean of the core community of motifs in each cluster. The core community was defined as the top 10% of motifs in each cluster with the most within-cluster nearest neighbors. Prior to averaging, motifs were aligned to a ‘template’ motif. The template motif shared the most zero-lag peak temporal cross-correlations with all other motifs. If there were multiple templates, one was chosen at random. All motifs were zero-padded to a length of *3L* (39 timepoints) and aligned to these templates by their maximal cross-correlation lag and then basis motifs calculated from the core communities. Basis motifs were then aligned to one another by shifting the center of mass of activity to the middle timepoint. Timepoints with no variance across all basis motifs were removed, resulting in basis motifs that were 26 timepoints (∼2s) long.

### Percent Explained Variance Calculations

For all experiments, the percent explained variance (PEV) in neural activity was defined as

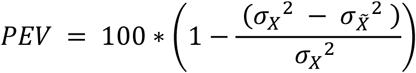

Where 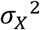 and 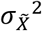 denote the spatio-temporal variance of the original data and reconstructed data respectively.

The PEV of individual motifs was calculated by convolving the motif with its temporal weighting and computing PEV as above (see *helper.reconstruct* function from *seqNMF* toolbox). Thus, the PEV of individual motifs reflected both the frequency of motif occurrence and the quality of fit to the data. Relative PEVs of individual motifs were calculated by dividing the PEV of each motif by the total PEV across all motifs for that epoch.

Timepoint-wise PEV (Fig. 3 supplement 1B) used the same calculation as above but was performed separately on each timepoint of an epoch. Thus, these analyses reflect solely the spatial variance in activity captured by reconstructed data for each individual timepoint.

### Cross-Temporal Autocorrelation Analysis

The cross-temporal autocorrelation of each motif was calculated by computing the spatial correlation between frames of the motif at vary temporal lags. The resulting autocorrelation was then fit with an exponential to estimate the half-life of decay in the autocorrelation (τ). Pixels with no variance across motif timepoints were ignored when calculating correlation.

### Static Networks

Static networks were generated for each motif by replacing all of the active timepoints of a motif with the mean activation of that motif (see Fig. 3E for example). Active timepoints were defined as any timepoints with variance across pixels greater than zero. Thus, static networks represent a constant ‘state’ of activation of the same brain regions for the duration of that motif. The temporal weightings of these static networks were refit to the withheld data the same way as dynamic motifs (i.e. the activation of these states could vary throughout an epoch; described above).

### Estimating the Working Resolution of Widefield Imaging

One concern is that the spatial resolution of our approach may have artificially limited the dimensionality of the observed motifs. Therefore, we sought to estimate the ‘working resolution’ of our approach by grouping pixels into functional clusters, defined as contiguous groups of pixels with correlated activity (Fig. 4 supplement 3).

To identify functional clusters, we divided each recording epoch into 1 second time periods (17,280 total 1-second periods, each with 13 frames). A pixelwise correlation matrix was computed for each 1-second period. Next, PCA was applied to each correlation matrix, producing a (pixel × component) matrix of ‘eigenconnectivities’. These eigenconnectivities reflected the dominant correlation patterns across pixels during that 1-second period (similar to approaches by Leonardi et al., 2013; Preti and Ville, 2017)

The first eigenconnectivity from each 1-second time period were concatenated together, creating a (pixel × 17,280) matrix that captured the wide variety of different possible spatial patterns across the cortex. We then used Phenograph (k=15) to group pixels according to their correlation across these spatial patterns. This produced 37 ‘functional clusters’ of highly correlated pixels (Fig. 4 supplement 3; 18 in left and 19 in right hemisphere. Similar results (∼20 clusters per hemisphere) were observed by performing PCA on each 1-second time period and then clustering as above (i.e. without first creating a correlation matrix; this creates an ‘eigenimage’ instead of an ‘eigenconnectivity’ as in Friston, 2004)

Several lines of evidence suggest our motifs were not constrained by the functional resolution of our approach. First, as shown in Figures 1, 3, and 4, most motifs are dynamic – different regions are activated over time. This suggests motifs are not limited by the spatial resolution of our approach, as the motifs capture the flow of activity across distinct spatial regions. Second, our imaging approach can still capture pixel-wise activity (Fig. 7B), suggesting the functional clustering is a lower-bound on our resolution. Even using this lower bound, the dimensionality of our motifs is far less than the possible dimensionality of our approach. For example, even if motifs engaged only 1-2 functional clusters, there are still 18^2^=324 possible patterns.

### Quantifying Spontaneous Behavioral States

Behavioral state was quantified using the binarized intensity of nose motion energy, whisker pad motion energy, and total limb speed. Nose and whisker pad motion energy was quantified as the mean absolute temporal derivative of pixels within a manually selected ROIs (shown on Fig. 5A). The whisking speed of the mouse was faster than the frame rate of the behavioral videos. This caused the whiskers to be blurred when the animal was intensely whisking, resulting in a low value for the whisker energy measure. Therefore, whisking motion energy was inverted for all analyses (e.g. 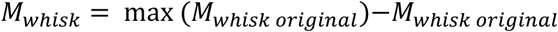) so that the axes were consistent across the three behavioral variables (i.e. higher values indicate higher energy).

Markerless tracking of paw position (DeepLabCut; Mathis et al., 2018) was used to measure limb speed. Training and validation of DeepLabCut neural network followed Nath et al.’s, 2019 published protocol, with the network trained to identify paw position as the center point between the 1^st^ and 5^th^ digit of each paw. To improve accuracy of limb position estimates, nose, tail base, and tail root were also tracked. 360 frames from 3 animals were used for network training. One refinement iteration was performed. The network was trained until loss plateaued (Fig. 5 supplement 1C, 120000 iterations). Total limb speed was calculated as the summed absolute temporal derivatives of the x and y position of all four paws.

To categorize behavioral state, a gaussian mixture model (GMM) was fit to the distributions of these three behavioral variables (MATLAB; *fitgmdist* function; 2 components). Timepoints were assigned to one of two behavioral states according to the 2-second running product of the posterior probability of being in each state. Any timepoints with a likelihood of being in either state that was less than 0.5 were excluded from analysis (this was less than 0.01% of all timepoints in both animals).

To decode behavioral state based on motif activity, we used a support vector machine classifier with radial basis function kernels (MATLAB; *fitsvm* function; Sequential Minimal Optimization solver). Prior to training a classifier, 20% of each trial type was held out as a validation test data set. For each classifier, two hyperparameters, “box-constraint” (a regularization parameter to prevent overfitting) and “kernel scale” were tuned using cross validation within the training set data (5 folds, balanced trial types: using the MATLAB functions *fitsvm* and *cvpartition*). Hyperparameters were optimized using Bayesian optimization (MATLAB, *bayesopt* function). Tuned classifiers were then tested on withheld validation data. Classifier accuracy was quantified using the area under the curve (AUC) of the receiver operator characteristic function (MATLAB, *perfcurve* function). Results of this classification analysis are reported in the main text.

### Fitting Average Traces to Sensory Trials

Average stimulus responses were calculated by taking the mean of all trials across animals for visual and tactile stimuli. Different stimuli within a modality were combined to generate the average trace (e.g. visual grating 1 and 2 were averaged together). Temporal weightings of average traces were refit to trials using the same seqNMF algorithm and parameters as when refitting basis motifs, except now the average stimulus response was used in place of the motifs. Additionally, prior to fitting, trials were zero-padded which allowed the algorithm to flexibility shift the average trace timing to best match the evoked response timing of each trial. This allowed us to directly compare the PEV of the motifs and the average stimulus response.

### Comparing Stimulus Evoked Motif Responses

We sought to compare the motif responses evoked by different stimuli (Fig. 6D-E). Motif activity was estimated by convolving each motif with its temporal weight on each trial (as described above; see *seqNMF Toolbox helper.reconstruct* function). Stimulus selectivity of a motif was estimated by comparing the evoked responses to each stimulus (two-sample t-test). This was done over time, using 300ms windows, stepped every 150ms to give a timecourse of similarity. Resulting p-values were Bonferroni corrected for multiple comparisons across time and conditions (i.e. 24 comparisons in Fig. 6D-E, 12 windows × 2 modalities).

### Pixelwise Classifier Analysis Comparing Visual Stimuli

As described above, mice were presented with two different visual stimuli: visual stimulus 1 (grating drifting from medial-to-lateral) and visual stimulus 2 (grating drifting from lateral-to-medial). Mice received 15 trials of each stimulus on 4-5 consecutive recording days (60-75 total of each type per mouse). As described in the main text, the response of the ‘visual’ motif (#10) did not differ between the two stimuli. To test whether this was due to a limitation in our imaging approach, we tested whether we could classify neural responses as belonging to the two stimuli (Fig. 7B)

To classify stimulation response, we used a support vector machine classifier (see detailed description above). Classification was performed on pixels restricted to the right hemisphere (contralateral to stimulus) and in the top 95% intensity percentile of motif 10 (visually-evoked motif). This resulted in a total of 6006 possible feature pixels (231 pixels across the 26 timepoints of visual stimulus presentation). For each classifier, we performed a cross-validated ANOVA feature selection to select a subset of these pixels as classification features (Pereira et al., 2009). The p-value for a one-way ANOVA comparing responses to each stimulus type was computed for each pixel in the training data. The 50 pixels with the highest -log10(p-values) were used for classification. After determining the feature vector, the classification hyperparameters were tuned within the training data, as described above.

For each mouse, pixelwise classification was performed on two datasets (Fig. 7B). First it was performed on the original data (after the preprocessing steps above). Second it was performed on the reconstructed activity of the motifs fitted to this original data. Thus, for both classification procedures, the data were the same spatial resolution. The only difference between the data sets were the residuals of the motif fitting procedure. Therefore, a total of 18 classifiers were trained (2 per animal).

### Evaluating Motif Expression During Two Tactile Stimuli

Animals were also presented with two different tactile stimuli; airpuffs that either traveled from anterior to posterior or from posterior to anterior. Motifs captured the vast majority of explainable variance in the response to tactile stimuli. In particular, Motif 1 (tactile-specific; Fig. 6E) captured a large portion of response to both stimuli (Fig. 6 supplement 1; 61.08% +/-1.97% SEM for non-stimulus specific motifs; 36.90% +/- 1.90% SEM for stimulus-specific motif 1).

However, it is important to note that, unlike the visual stimuli, these two tactile stimuli were likely significantly different in their impact on behavior. First, the behavioral connotation of an anterior-approaching stimulus is likely different than that of a posterior stimulus. Second, although both airpuffs were directed at the whisker pad, differences in the solenoids used to control the airflow impacted flow rate and air pressure. Thus, one may expect these stimuli to evoke different intensity responses (and thus different motif temporal weightings). Consistent with this hypothesis, the relative percent variance in neural activity captured by motifs differed in response to the two airpuffs (Fig. 7 supplement 1A). Because of these large differences at the level of motifs, there was no need for to classify the tactile stimuli. However, stimulus-specific activity in the residuals still only captured a small part of the total explainable variance (2.02% +/- 0.38% SEM; Fig. 7 supplement 1B). Thus, as we saw for visual stimuli, most cortex-wide neural activity can be explained by the expression of motifs.

## Supplemental Information

**Supplemental Movie 1. Example Widefield Imaging of Cortical Activity.** Shows an example of widefield imaging (left) alongside a video of animal’s behavior (right).

**Supplemental Movie 2. Example Fit of Data Reconstructed from Discovered Motifs. (A)** Temporal weightings of discovered motifs. **(B)** Original data. Color indicates fluorescence intensity of original data (i.e. 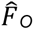: normalized between 0 to 1, using the 99.9^th^ percentile of all pixels). **(C)** Reconstructed data. Created by convolving motifs with their temporal weightings (see Method for details). Color indicates intensity of reconstructed data (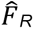); color scale follows **B**. **(D)** Residual of fit (i.e. 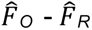). Color scale indicates the quality of fit of reconstructed to original data. Red dots indicate bregma.

**Supplemental Movie 3. All Basis Motifs.** Basis motif display follows Figure 4C. For video visualization, all motifs were convolved with a 450ms gaussian temporal weighting vector to mimic a ‘spike’ in activity.

**Supplemental Table 1.**
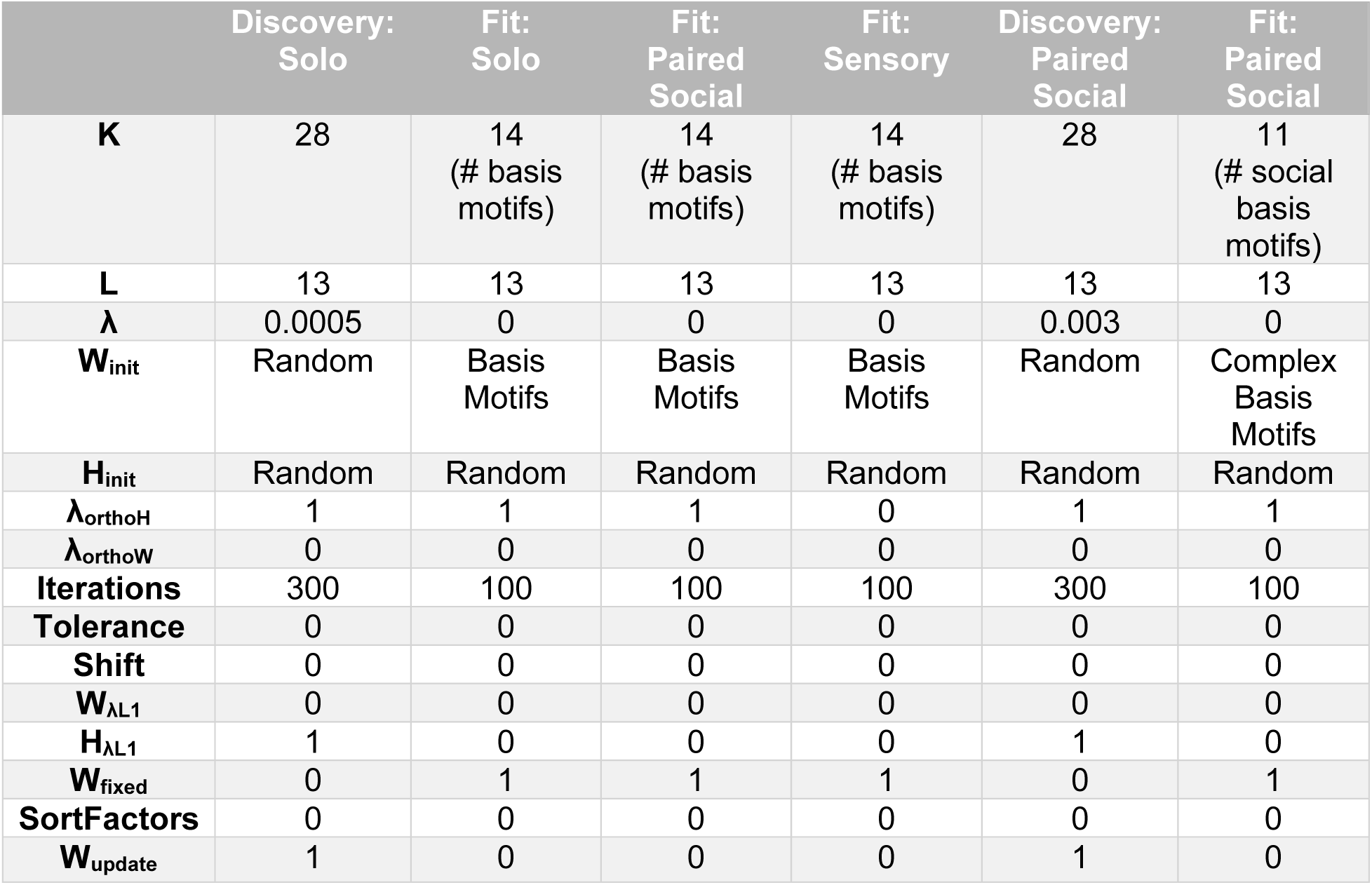
CNMF parameters used in each experiment. See methods for descriptions of parameter choices and fitting procedures. For complete details refer to seqNMF MATLAB Toolbox (Mackevicius et al., 2019).

**Figure 1, Supplement 1.**
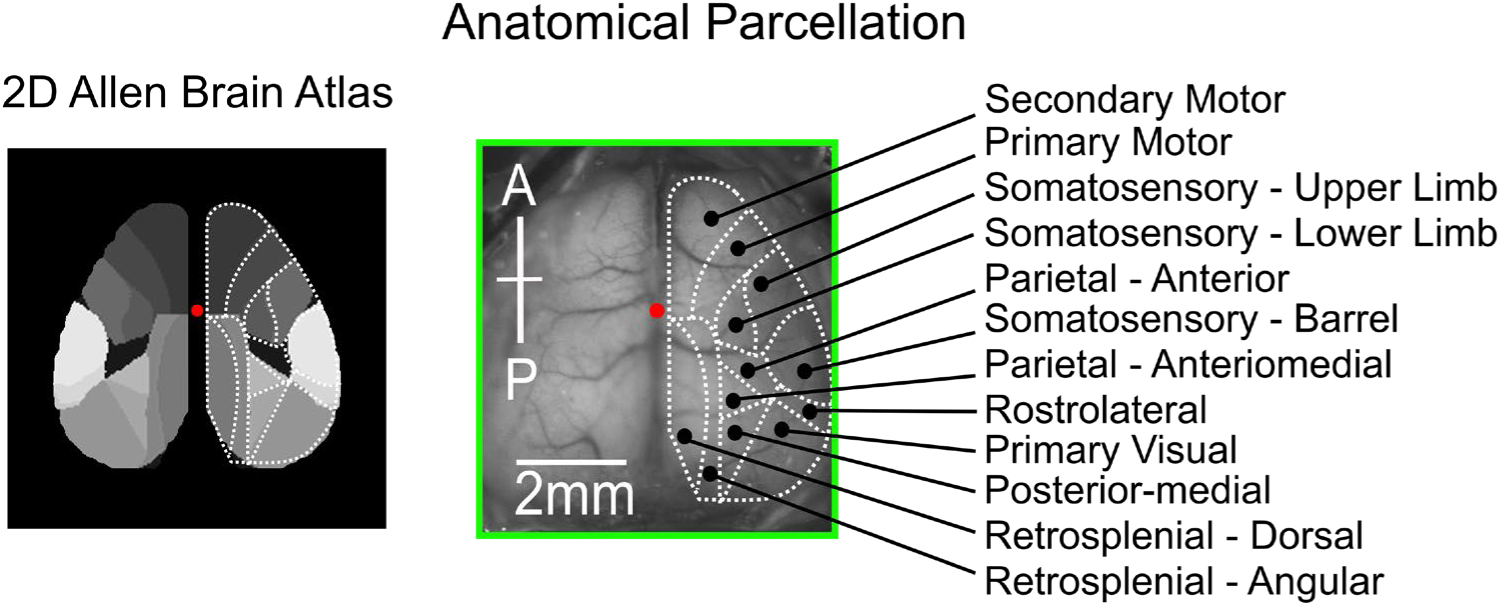
Detailed anatomical parcellation. **Left:** 2D projection of Allen Brain Atlas anatomical parcelation. **Right:** Allen Brain Atlas anatomical region labels overlaid on example mouse brain. Dotted white lines indicate manually drawn region outlines overlayed on motifs in main figure text.

**Figure 1, Supplement 2.**
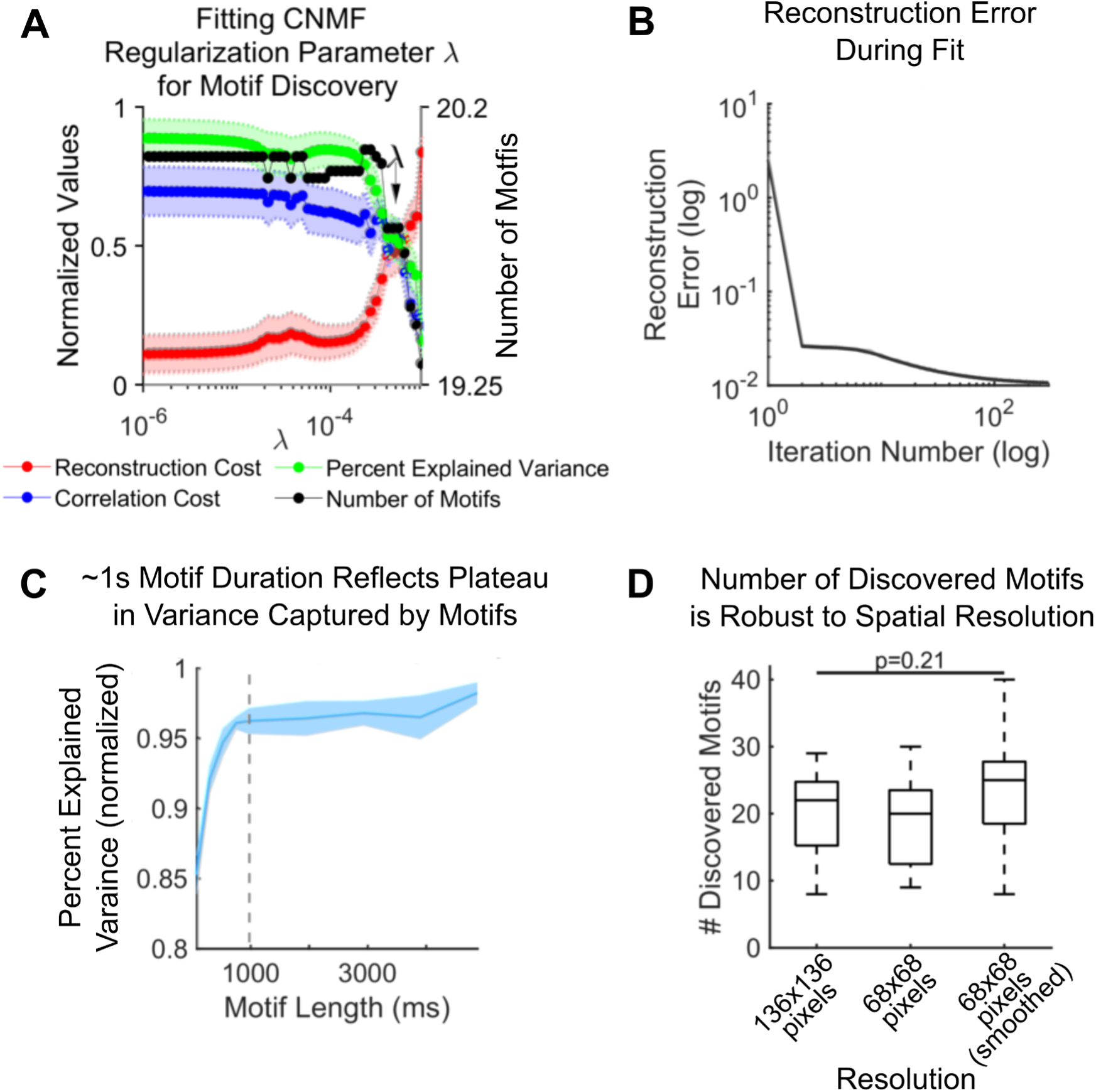
Testing impact of CNMF parameters on motif discovery. **(A)** The effect of changing the spatio-temporal regulation paramater (λ) in the CNMF algorithm on reconstruction cost (red), correlation cost (blue), explained variance (green) and number of identified motifs (black). Each data point indicates the mean value from 20 fit epochs (randomly selected; no replacement); shaded regions indicate SEM. Y-axis units are arbitrary; values were normalized between 0 and 1 across λ values for each of the 20 fits. Chosen lamda is indicated by arrow. **(B)** Reconstruction error as a function of iteration number of CNMF algorithm. All motif discovery factorizations were run for 300 iterations, at which point there was minimal improvement in reconstruction error. **(C)** Post-hoc validation of choice of motif duration. Motifs were discovered using motif durations up to 5 seconds. 1 second motifs captured comparable variance to longer motif durations. Shaded regions and dark line show mean and SEM, respectively. **(D)** Spatial resolution did not change number of discovered motifs. Spatial resolution indicated along x-axis. When smoothed, a 2D gaussian filter (σ = [1,1] pixels; see Methods for details) was convolved across each frame. Line, box, and whiskers denote median, 25^th^-75^th^ percentile, and range repectively. Significance estimated with one-way ANOVA.

**Figure 3, Supplement 1.**
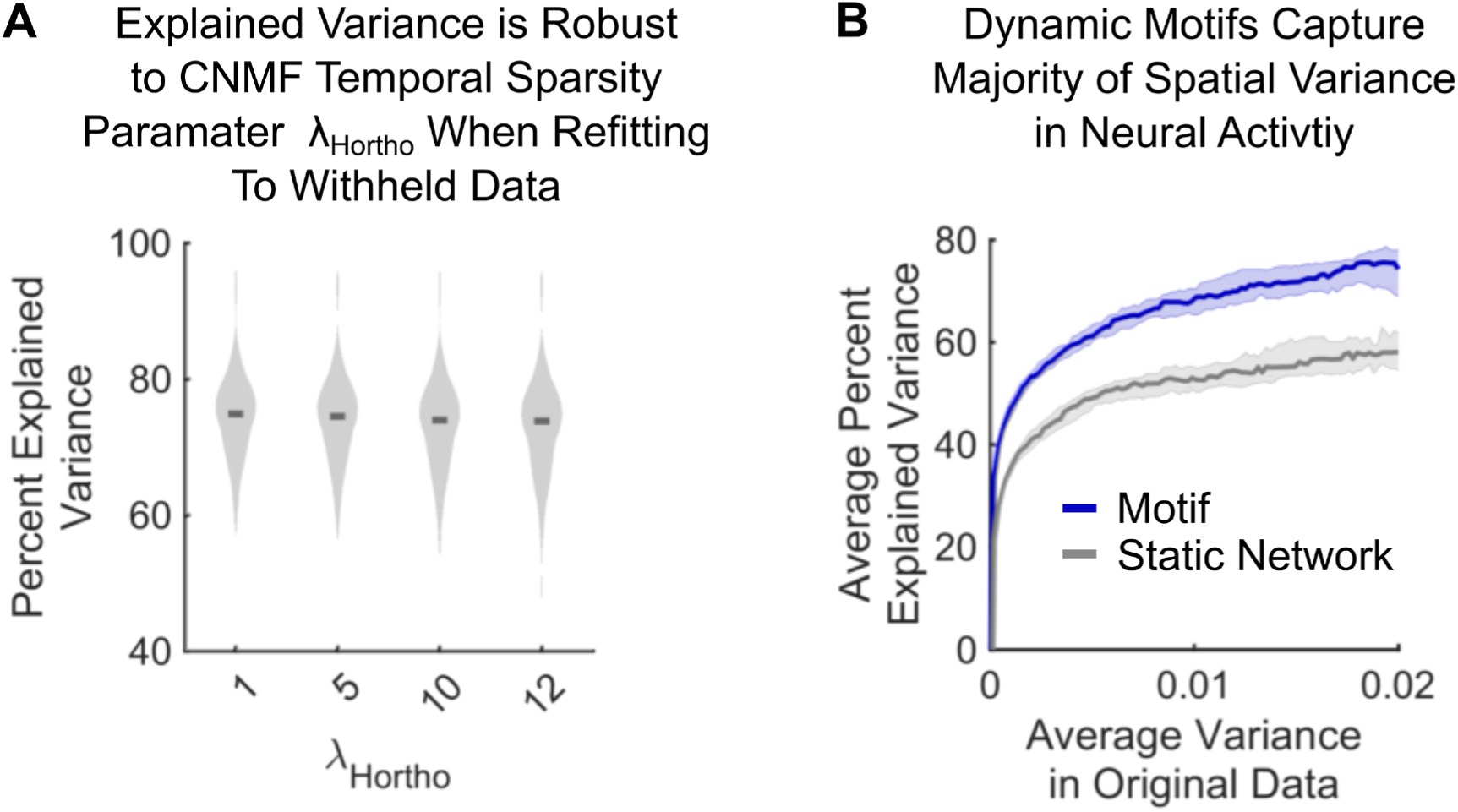
**(A)** Percent of variance in neural activity explained by motif reconstructions as a function of temporal sparsity parameter λ_Hortho_. Full distribution shown. Dark lines indicate median. **(B)** Percent of variance in neural activity explained by motif reconstructions (purple) and static networks (gray) of withheld epochs per timepoint. The explained variance was separately calculated for each timepoint of each epoch (1560 timepoints per epoch, 144 epochs; see Methods for details). Timepoints were then binned according to the variance across the image in the original data (100 equal bins). The explained variance captured by the reconstruction was averaged per bin per epoch. Dark lines indicate median of 144 epochs. Shading indicates 95% confidence interval. Analyses performed on withheld data within the same animals (as in Fig. 3D; purple).

**Figure 4, Supplement 1.**
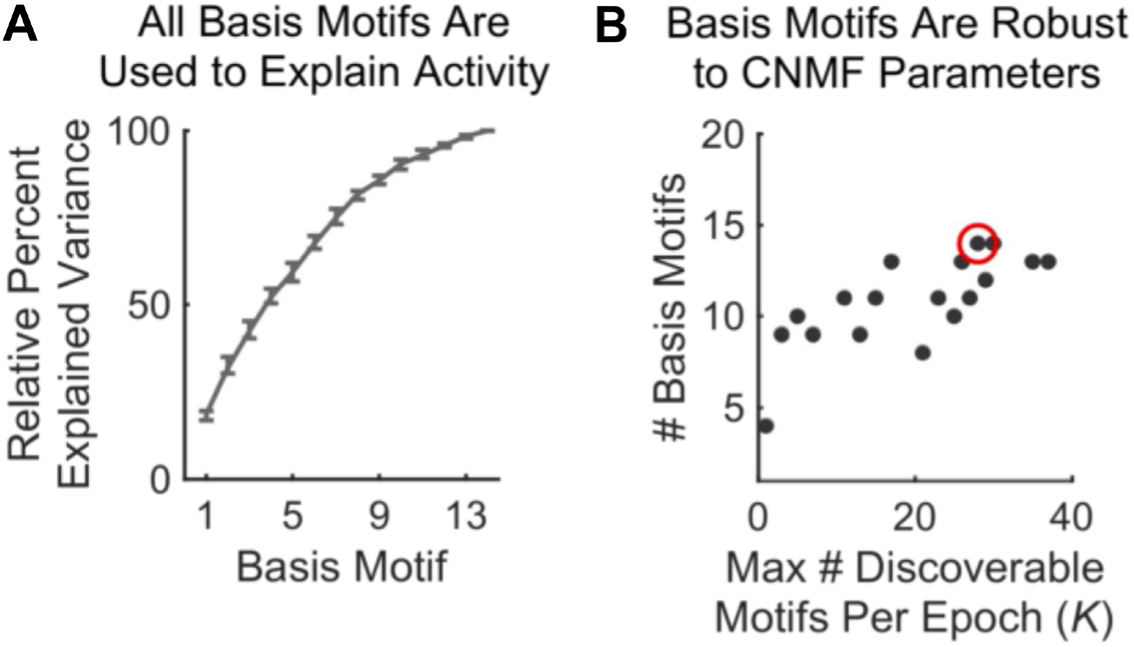
Additional characterizations of basis motifs. **(A)** Cumulative sum of relative percent explained variance of basis motifs. Relative PEV was defined as the fraction of total PEV (of all motifs) captured by each motif per epoch. Basis motifs are in descending order by their relative PEV; these labels are used for basis motifs throughout manuscript. Line and error bars denote mean and 95% CI, respectively. **(B)** Number of basis motifs discovered (y-axis) as a function of CNMF hyperparameter K, the maximum number of discoverable motifs allowed in a single epoch (x-axis). Motif discovery and clustering was repeated for each K value (see Methods for details). Regardless of parameters, 10-14 basis motifs were identified. Red circle denotes K value (28) used for all experiments in the main text, conservatively chosen to maximize the number of basis motifs discovered.

**Figure 4, Supplement 2.**
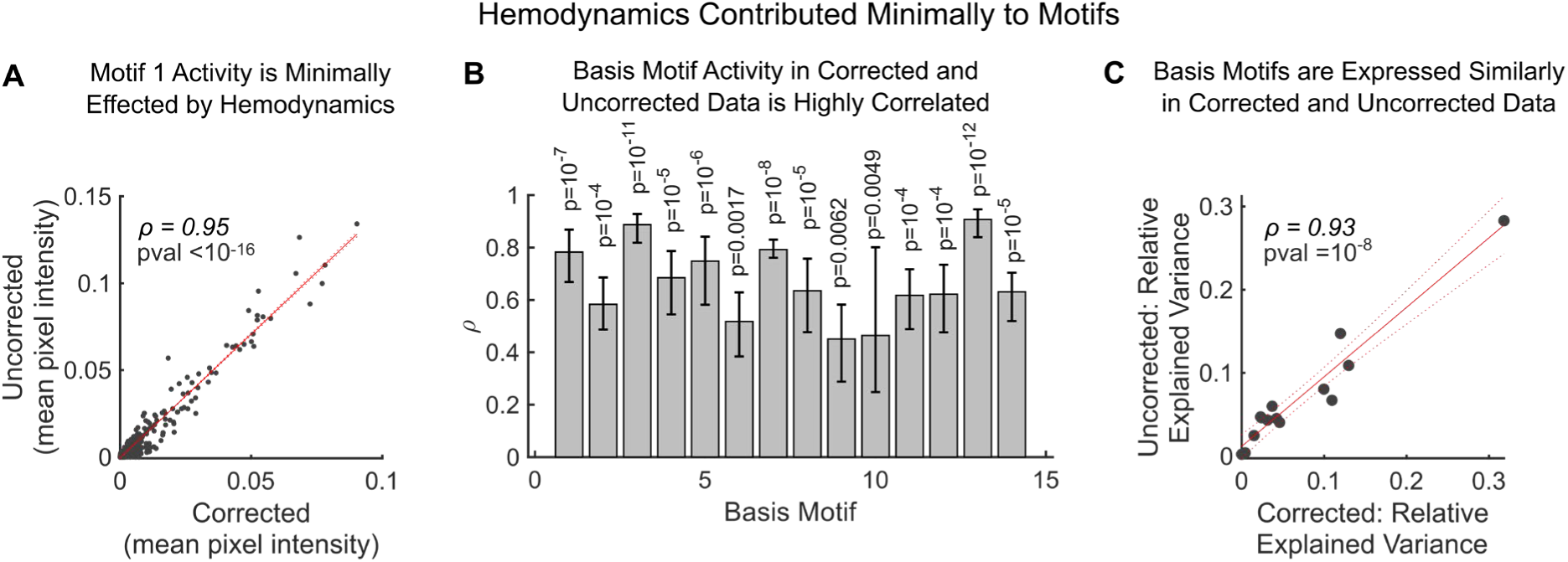
Multiwavelength hemodynamic correction. **(A)** Example correlation between average pixel intensity of motif 1 reconstruction before and after multiwavelength hemodynamic correction (see Methods for details). Gray markers represent mean pixel intensity per timepoint. Solid and dotted red lines show linear least squares fit and 95% confidence bounds respectively. **(B)** The correlation in activity between corrected and uncorrected data was high for all motifs. Correlation is shown for N=30 2-min epochs across 2 animals. Mean and confidence intervals calculated on fisher z-transformed data before reconverting to Pearsons correlation coefficient. **(C)** Average relative variance explained by motifs in corrected and uncorrected epochs. Data points show mean of N=30 2-min epochs. Display follows **A**.

**Figure 4, Supplement 3.**
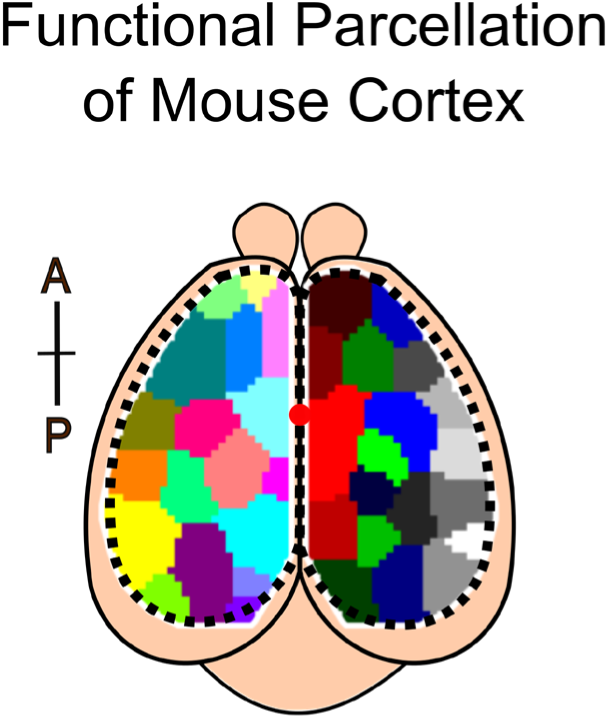
Estimating the ‘working resolution’ of widefield imaging approach. Parcellation of mouse cortex into functional clusters (N = 18 and 19 for left and right hemisphere, respectively). Functional clusters grouped pixels that were correlated over time (see Methods for details). Each color denotes a separate functional cluster. Red dot indicates bregma.

**Figure 4, Supplement 4.**
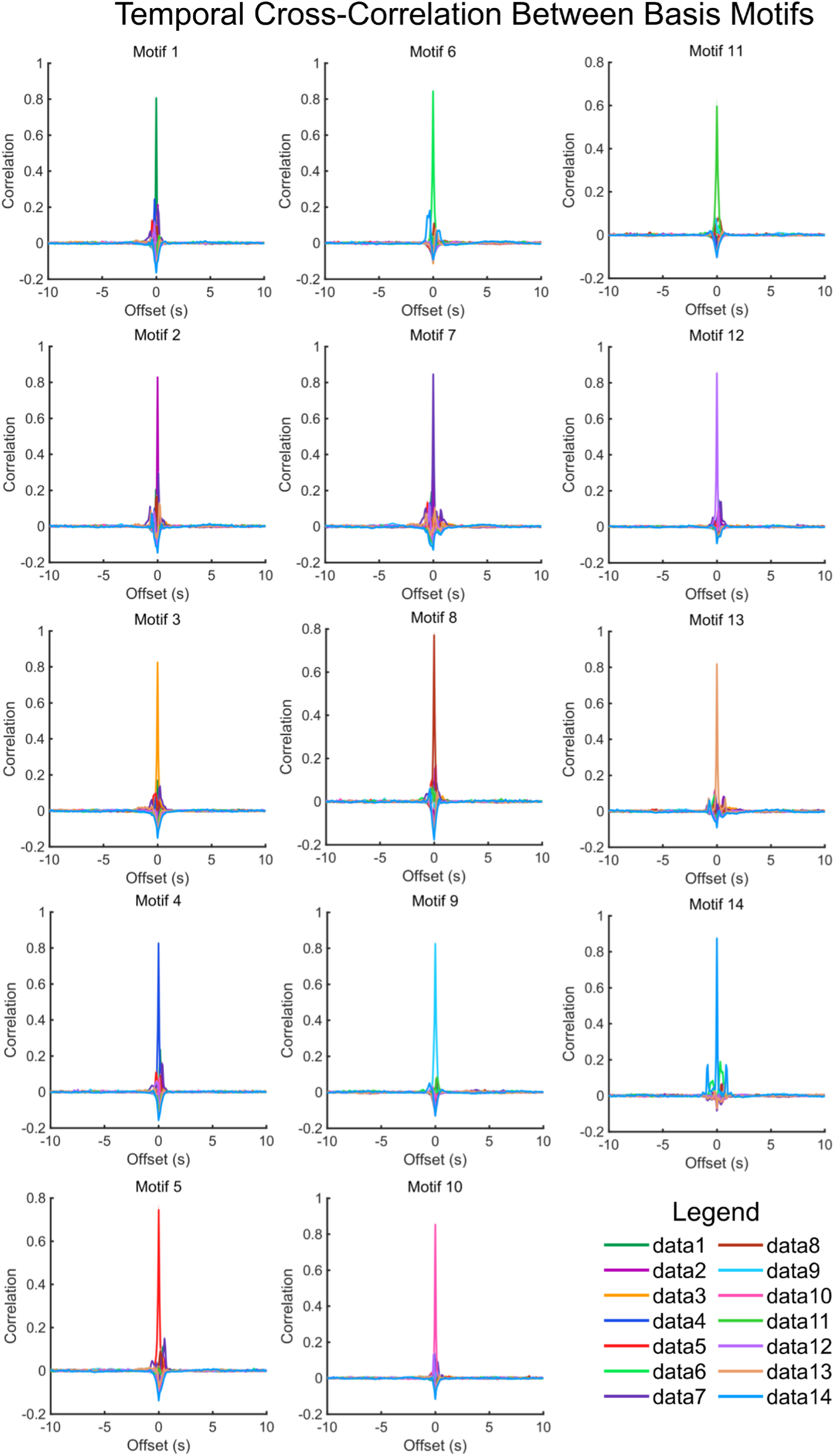
Temporal cross correlation between motifs. Temporal cross-correlations (and autocorrelations) performed on the temporal weightings of basis motifs refit to N=144 withheld epochs. Line and shading reflect mean and SEM respectively. No obvious hierarchical structure was observed in the activation of different motifs.

**Figure 7, Supplement 1.**
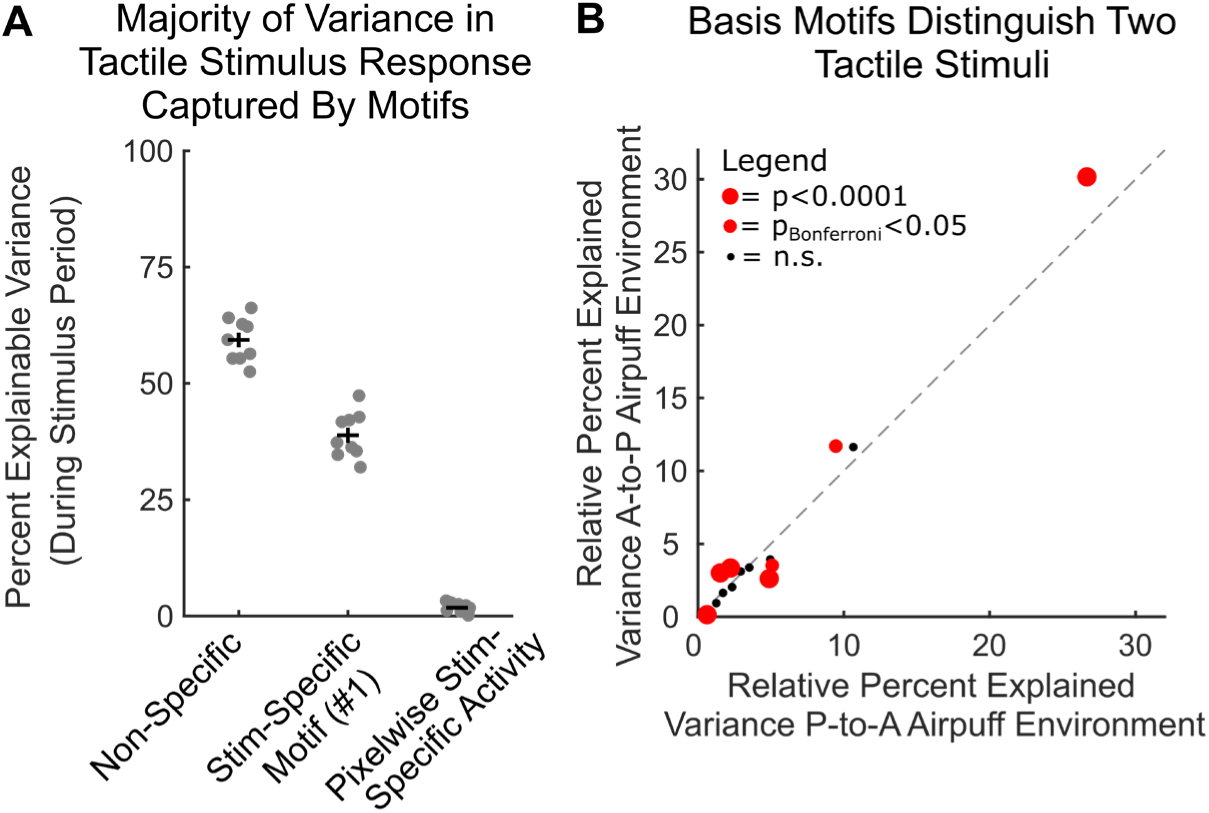
Basis motif expression differs in response to two tactile stimuli. **(A)** The majority of variance in neural activity could be explained by motif activity, not stimulus-specific activation. Plot shows the percent of explainable variance in the neural response to tactile stimuli that is captured by non-specific motifs (left column), the stimulus specific motif (motif 1; middle column) and stimulus-specific residuals (right column). Follows Figure 7C. Data points correspond to mice (N=9). Black horizontal bars indicate mean and vertical bars indicate SEM. **(B)** Plot shows the relative percent explained variance for each basis motif in response to anterior-to-posterior airpuffs and a posterior-to-anterior airpuffs directed at the whisker pad. Presentation follows Figure 6B.

